# Neural dynamics express syntax in the time domain during natural story listening

**DOI:** 10.1101/2024.03.19.585683

**Authors:** Cas W. Coopmans, Helen de Hoop, Filiz Tezcan, Peter Hagoort, Andrea E. Martin

**Affiliations:** Max Planck Institute for Psycholinguistics, Nijmegen, The Netherlands; Donders Institute for Brain, Cognition and Behaviour, Radboud University, Nijmegen, The Netherlands; Centre for Language Studies, Radboud University, Nijmegen, The Netherlands

**Keywords:** hierarchy, syntax, language comprehension, magnetoencephalography, temporal response function

## Abstract

Studies of perception have long shown that the brain adds information to its sensory analysis of the physical environment. A touchstone example for humans is language use: to comprehend a physical signal like speech, the brain must add linguistic knowledge, including syntax. Yet, syntactic rules and representations are atemporal (i.e., abstract and not bound by time), so they must be translated into time-varying signals for speech comprehension and production. Here, we test three different models of the temporal spell-out of syntactic structure against brain activity of people listening to Dutch stories: an integratory bottom-up parser, a predictive top-down parser, and a mildly predictive left-corner parser. These models build exactly the same structure but differ in when syntactic information is added by the brain – this difference is captured in the (temporal distribution of the) complexity metric ‘incremental node count’. Using temporal response function models with both acoustic and information-theoretic control predictors, node counts were regressed against source-reconstructed delta-band activity acquired with magnetoencephalography. Neural dynamics in left frontal and temporal regions most strongly reflect node counts derived by the top-down method, which postulates syntax early in time, suggesting that predictive structure building is an important component of Dutch sentence comprehension. The absence of strong effects of the left-corner model further suggests that its mildly predictive strategy does not represent Dutch language comprehension well, in contrast to what has been found for English. Understanding when the brain projects its knowledge of syntax onto speech, and whether this is done in language-specific ways, will inform and constrain the development of mechanistic models of syntactic-structure building in the brain.

## 1. Introduction

How the brain transforms continuous sensory stimulation into cognitive representations is a major question in human biology. Because sensory input alone is typically insufficient to uniquely determine a coherent and reliable percept, perception requires the brain to add information to its analysis of the physical environment (Wei & Körding, 2011). Language processing is a prime example: one must know a language in order to perceive it from speech, so comprehending spoken language requires the encoding of information beyond what is presented in the raw speech signal. This includes the formation of syntactic structures – the ‘instructions’ by which words hierarchically are combined for interpretation. What is noteworthy about this process is that (the brain’s) representations of syntactic rules and structures are abstract and not bound by time, yet they must be translated into temporal signals to comprehend and produce speech. Here, we compare three competing ways in which syntactic information can be metered out by the brain. These different models of parsing build exactly the same syntactic structure, but differ in the dynamics of structure building. We thus investigate not only what kind of linguistic information the brain projects onto its perceptual analysis of speech, but also when it does so.

As this discussion makes clear, integrating the computational level of linguistic theory with the implementational level of cognitive neuroscience requires a linking hypothesis that specifies a connection between two fundamentally different types of data – atemporal linguistic structure and continuously varying neural dynamics (Embick & Poeppel, 2015; Poeppel, 2012). When it comes to studying syntactic structure building in the human brain, it is therefore important to be explicit about the structure of the syntactic representations, the algorithmic procedures for computing these representations in real time, and the linking theory that maps the output of these algorithms onto neural signals (Coopmans & Zaccarella, 2023; Demberg & Keller, 2019; Martin, 2016, 2020; Martin & Doumas, 2017, 2019; Sprouse & Hornstein, 2016). One promising approach relies on neuro-computational language models, which are computationally precise models of language processing that define a measure of processing difficulty for every word in a dataset that reflects everyday language use (Brennan, 2016; Hale et al., 2022). By determining whether this measure reliably predicts brain activity elicited by the words in that dataset, we can establish whether there is evidence for the hypothesized linguistic computations in the neural signal.

In the current study, we compare different neuro-computational models in terms of their ability to predict brain activity of people listening to Dutch audiobook stories. In order to connect syntactic structure with neural dynamics, we adopt the complexity metric ‘incremental node count’, which corresponds to the amount of structure that must be built when incorporating a word into the hierarchical structure of the sentence. Incremental node count is an appropriate linking hypothesis because it both directly reflects atemporal syntactic structure and also varies in time. By expressing syntax in the time domain, the dynamics of incremental node count model the temporal distribution of perceptual cues for the inference of syntax (Figure 1). Here, we regress node count against brain activity in a time-resolved manner, in order to uncover the neural dynamics of syntactic structure building during natural story listening.

**Figure 1.**
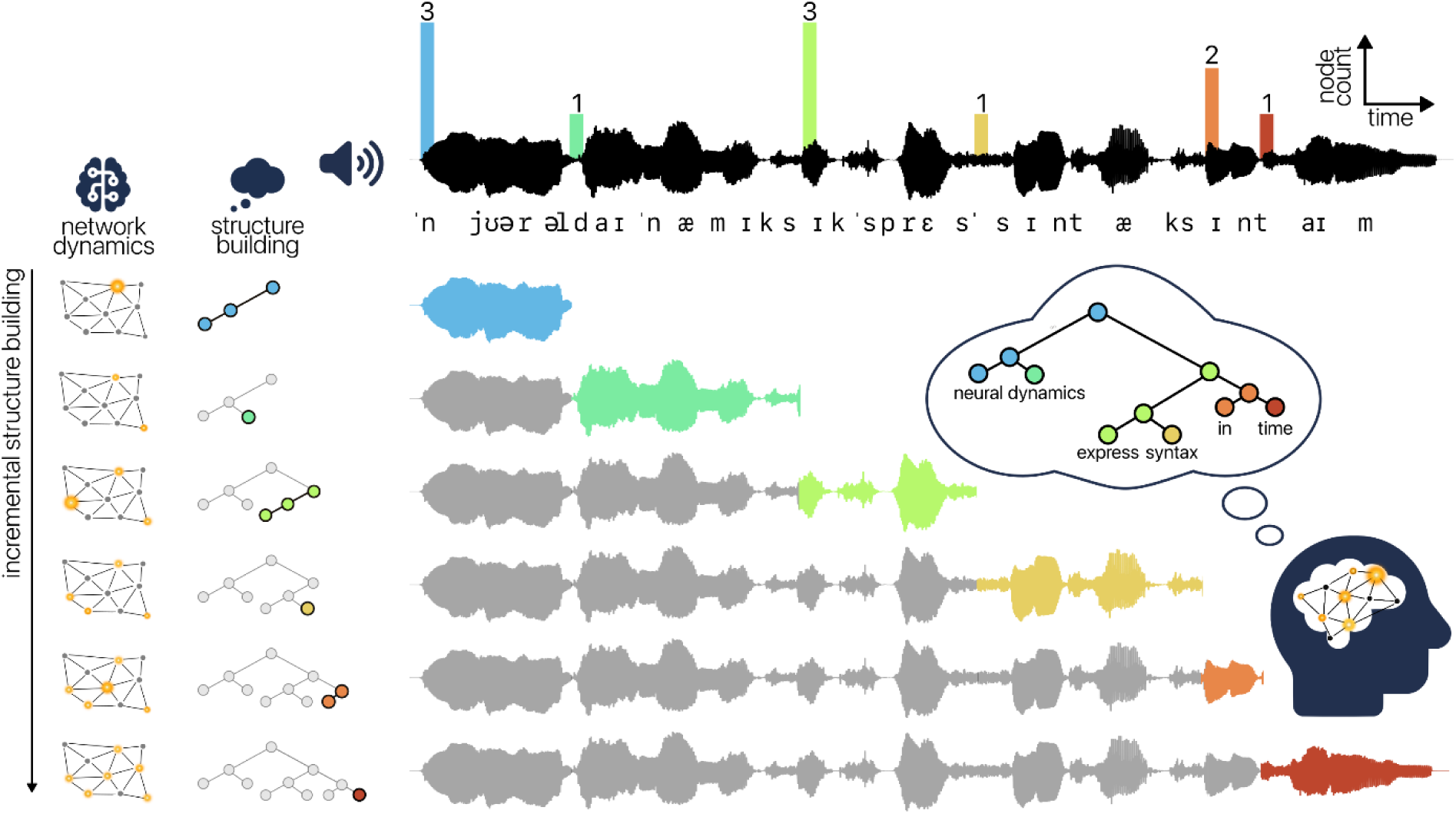
Neural dynamics of incremental structure building in spoken language comprehension. Speech is segmented into words, which are incrementally integrated into a hierarchical syntactic structure with a meaningful interpretation. The amount of structure built at each word (incremental node count) is reflected in the dynamics of the network, which lead to a neural signal measurable with magnetoencephalography.

### 1.1 Neuro-computational models of sentence comprehension

Brennan (2016) defines neuro-computational models as consisting of a parser *P_G,A,O_* that contains a grammar *G* with rules to construct representations, an algorithm *A* for incrementally applying the grammar word by word, and an oracle *O* that resolves indeterminacies. When applied to a sequence of words *w_1_,w_2_,…w_n_*, *P_G,A,O_* yields a sequence of mental states *m_1_,m_2_,…m_n_*, which correspond to (partial) syntactic structures. These mental states can be quantified via an auxiliary hypothesis or linking rule, often referred to as complexity metric in psycholinguistics because of the way in which it quantifies language processing complexity (Hale et al., 2018). The complexity metric *C* thus represents the neural state in quantifiable cognitive terms; it stands for estimated brain states. The estimated brain states are linked to the observable neural signal via a response function *R*.

This neuro-computational method has the potential to evaluate the cognitive and neural relevance of linguistic constructs to the extent that there are appropriate linking hypotheses about how these constructs are manifested in neural activity. One assumption underlying this approach is that a model that captures linguistic competence give rises to measures that are predictive of experimental data. It is promising that the results of recent neuroimaging studies point in this direction. For instance, grammars that compute hierarchical structure account for variance in neural activity above and beyond the variance accounted for by sequence-based models (Brennan et al., 2012, 2020; Brennan & Hale, 2019; Lopopolo et al., 2021; Martin & Doumas, 2017; Shain et al., 2020), and hierarchical grammars that naturally represent long-distance dependencies, which are ubiquitous in natural languages, uniquely predict activity in brain areas commonly linked to syntactic structure building (Brennan et al., 2016; Li & Hale, 2019; Nelson et al., 2017; Stanojević et al., 2023). These findings reinforce the view that grammars that are well-equipped to account for natural language structures (competence) are also required to adequately model the activity of the brain when it incrementally computes these structures (performance).

#### 1.1.1 Three models of the temporal spell-out of syntactic structure

Parsing models specify how syntactic parse states incrementally unfold in time during language comprehension. In the current work, these parse states correspond to partial syntactic structures, which are generated via the rules of X-bar theory (the grammar *G*; Carnie, 2021; Jackendoff, 1977). We chose to construct X-bar tree structures because they are appropriately expressive to deal with natural language structures (e.g., long-distance dependencies, movement) and because recent work using neuro-computational models has shown that complexity metrics derived from these structures predict brain activity above and beyond activity accounted for by both sequential models and models that are hierarchical but less expressive (Brennan et al., 2016; Li & Hale, 2019; Nelson et al., 2017). We further assume a perfect oracle *O*, which resolves temporary ambiguities in the right way (Bhattasali et al., 2019; Brennan et al., 2012, 2016; Brennan & Pylkkänen, 2017). This means that the parser builds the correct structure at any point in the parse, even when faced with locally ambiguous input.

Any given syntactic structure may be built in different ways, depending on the parsing algorithm *A* that is adopted. We will be concerned with three algorithms for building structure: a top-down algorithm, a bottom-up algorithm and a left-corner algorithm (see Hale, 2014).^1^ The top-down parsing method works via *expansion* of rewrite rules. Before each word, all rules necessary to attach the upcoming word to the structure are expanded (e.g., in VP → V NP, the VP is expanded as V and NP). As these rules are applied based on the left-hand side of the rule and in advance of each word, this method builds constituent structure entirely predictively. The bottom-up parsing method, instead, builds constituent structure in a non-predictive manner, as it postulates a constituent node only after all of its daughter nodes are available. This process is referred to as *reduction*: when all information on the right-hand side of a rule is available, the input is reduced to the constituent node (e.g., in VP → V NP, the input V and NP is reduced to VP). In between these two parsing methods is a mildly predictive left-corner strategy, which works via *projection*. A constituent node is projected after the very first symbol on the right-hand side of the rewrite rule (its left corner) is seen (e.g., the VP node is projected when V is available). This strategy is only mildly predictive because, while it requires input to build structure (in contrast to the top-down strategy), the input can be incomplete (in contrast the bottom-up strategy).

To illustrate how these strategies differ in temporal spell-out, consider a simple sentence like ‘the boy sleeps’. The total number of operations (`expand`, `reduce`, `project`) for the three parsing methods is the same, but the timepoints at which they are applied differ. The predictive top-down parsing strategy postulates structure early in time. For instance, three operations are applied before ‘the’, corresponding to expansion of the S, NP, and D nodes (Table 1). Only one operation is applied at ‘the’ on the bottom-up method, because there is complete evidence for the determiner node D only. The next word ‘boy’ is the second word of the noun phrase, so two operations are applied bottom-up, but only one is applied top-down. What this simple structure illustrates is that these parsing methods differ in the dynamics of structure building. They make different claims about the time points at which processing complexity is high, and these differences will only be magnified when the sentences become longer and more complex.

**Table 1.** Parser actions of top-down, bottom-up and left-corner parsers for the sentence ‘the boy sleeps’. The `scan/shift` action corresponds to processing or moving to the next word in the sentence and is not a structural operation. The other operations are explained in the text.

| Top-down |  | Bottom-up |  | Left-corner |  |
| --- | --- | --- | --- | --- | --- |
| expand by | $S \rightarrow NP VP$ | shift | the | shift | the |
| expand by | $NP \rightarrow D N$ | reduce by | $D \rightarrow the$ | project | $D \rightarrow the$ |
| expand by | $D \rightarrow the$ | shift | boy | project | $NP \rightarrow D N$ |
| scan | the | reduce by | $N \rightarrow boy$ | shift | boy |
| expand by | $N \rightarrow boy$ | reduce by | $NP \rightarrow D N$ | project | $N \rightarrow boy$ |
| scan | boy | shift | sleeps | project | $S \rightarrow NP VP$ |
| expand by | $VP \rightarrow V$ | reduce by | $V \rightarrow sleeps$ | shift | sleeps |
| expand by | $V \rightarrow sleeps$ | reduce by | $VP \rightarrow V$ | project | $V \rightarrow sleeps$ |
| scan | sleeps | reduce by | $S \rightarrow NP VP$ | project | $VP \rightarrow V$ |

Here, we represent the number of parser actions at each word in the form of incremental node count, which corresponds to the number of new nodes in a partial syntactic structure that are visited by the parser when incrementally integrating a word into the structure (complexity metric *C*; Brennan et al., 2012; Frazier, 1985; Miller & Chomsky, 1963). Depending on the parsing algorithm that is used, node count reflects the number of `expand` (top-down), `reduce` (bottom-up) or `project` (left-corner) actions between successive words (Table 1), all of which can be taken as roughly corresponding to the syntactic load or complexity of those words. Previous neuroimaging work has shown that node count effectively quantifies syntactic complexity in cognitive terms (Bai et al., 2022; Bhattasali et al., 2019; Brennan & Martin, 2020; Brennan & Pylkkänen, 2017; Brennan et al., 2012, 2016; Coopmans et al., 2022; Giglio et al., 2024; Li & Hale, 2019; Lopopolo et al., 2021).

The left-corner strategy is thought to be cognitively plausible as a model of human language processing. It correctly predicts processing difficulty for center-embedded constructions (Abney & Johnson, 1991; Johnson-Laird, 1983; Resnik, 1992), is compatible with a range of findings from the sentence processing literature (Hale, 2014), and accounts for brain activity during language processing (Brennan & Pylkkänen, 2017; Nelson et al., 2017). However, much of the relevant psycholinguistic work has been done in English. A possible reason for the observation that left-corner parsing works well for English is that English phrases are strictly head-initial. The left corner of a phrase will therefore most often be its head, which can be used to build structure predictively (Arai & Keller, 2013; Boland & Blodgett, 2006; Schütze & Gibson, 1999). As English has certain grammatical properties that make it particularly well-suited for left-corner parsing, these findings need not generalize to (the processing of) other languages. Indeed, it is quite likely that people’s parsing strategies depend on the properties of the linguistic input, including the grammatical properties of the language in question. We therefore compare different parsing methods in terms of their ability to model syntactic processing of Dutch. In contrast to English, Dutch exhibits mixed headedness, with the verbal projections VP and IP being head-final. The left corner of a Dutch head-final VP will often be a multi-word constituent, such as in sentence (1) below:

1. De student heeft een essay over syntaxis geschreven. The student has an essay on syntax written ‘The student has written an essay on syntax.’

Notice the difference in word order between the Dutch example and its English translation. In English, the main verb precedes its complement (‘written – an essay on syntax’), thus yielding the head-initial order that is characteristic of English phrases. The reverse order is found in Dutch, where the verb follows its complement (*een essay over syntax* – *geschreven*, ‘an essay on syntax – written’), giving the head-final order. The left-corner method predicts that the VP constituent in Dutch will be projected only after the entire preverbal NP complement has been processed. This is unrealistically late, in particular if speakers of head-final languages adopt incremental or even predictive parsing strategies (Coopmans & Schoenmakers, 2020; Vasishth et al., 2010). It might thus very well be that left-corner parsing is not the best strategy for Dutch structures. Dutch is typologically related to English, but the head-finality of its verb phrases might invite different processing strategies within the neural language network (Bornkessel-Schlesewsky & Schlesewsky, 2016).

Note that our use of three different parsing methods does not imply that the human brain hosts multiple parsers. We use these methods as explicit and interpretable tools to derive values for incremental node count, a variable that quantifies the operations involved in syntactic structure building (e.g., the parser actions in Table 1). Thus, we use node counts as time-sensitive proxies for the operations that build structure in integratory or predictive ways, depending on how nodes (or, rather, parser actions) are counted. When node count derived from the top-down method significantly predicts brain activity, it does not mean that the brain is actually building hierarchical structure from the top of the tree to the bottom. Rather, it suggests that Dutch sentence comprehension can be characterized via a predictive mechanism, which postulates structure early in time. If node counts derived from the bottom-up and left-corner methods additionally explain variance in brain activity, it means that hierarchical structure is also built in integratory and mildly predictive manners, respectively. Importantly, these strategies need not be mutually exclusive, and might even reflect one and the same mechanism. What makes them different is the completeness of the input they require to build structure, with the top-down method building structure maximally eagerly (using uncertain and incomplete input), while the bottom-up method requires complete input and is therefore minimally eager (Stanojević et al., 2023). As different sentences and sentence positions vary in the extent to which they allow for predictive structure building, the comprehension process in full might thus be characterized in terms of multiple different processing strategies.

#### 1.1.2 From cognitive to brain states via temporal response functions

The complexity metrics derived from different parsing models are regressed against electrophysiological brain activity through multivariate temporal response functions (mTRFs). TRFs are linear kernels that describe how the brain responds to a representation of a (linguistic) feature (Brodbeck et al., 2018, 2022). This approach is similar to that of recent neuro-computational fMRI studies, which use the canonical hemodynamic response function to fit syntactic predictors onto brain activity in a given region of interest (Bhattasali et al., 2019; Brennan et al., 2012, 2016; Giglio et al., 2024; Li & Hale, 2019). But rather than assuming the shape of the response function, with the TRF method a response function can be estimated for each predictor separately, thus supporting time-resolved analyses. Moreover, by using multivariate TRFs, the acoustic properties of the auditory stimulus can be explicitly modeled, which is important for two main reasons. First, high-level linguistic features can be correlated with low-level stimulus properties, such that neural effects attributed to linguistic processing can also be explained as the brain’s response to non-linguistic, acoustic information (Daube et al., 2019). And second, because the neural response to acoustic properties is orders of magnitude larger than that to linguistic features (Brodbeck et al., 2022; Gillis et al., 2021; Tezcan et al., 2022), acoustic variance could mask subtle effects of linguistic information that are hiding in the data (Coopmans et al., 2022). Such low-level factors have not always been modeled in previous work using neuro-computational models of syntactic structure building, limiting their interpretability (e.g., Brennan & Hale, 2019; Lopopolo et al., 2021). Conversely, TRF studies commonly analyze predictors that reflect lexical information at most; they are rarely extended to capture super-lexical features. To address these two limitations of previous work, the current study uses the TRF method to evaluate the brain’s sensitivity to syntactic information in natural speech with high temporal precision, while appropriately controlling for lower-level factors.

### 1.2 The present study

In the current study, we compare different neuro-computational models in terms of their ability to predict source-reconstructed MEG activity of people listening to Dutch audiobook stories. The three models we evaluate rely on the same grammatical assumptions (X-bar theory), linking hypothesis (node count), and type of response function (TRF), but they differ in the parsing algorithm by which they build syntactic structure, and thus in the way they express syntax in the time domain. Based on recent results linking syntactic processing to delta-band activity (Bai et al., 2022; Brennan & Martin, 2020; Coopmans et al., 2022; Ding et al., 2016; Kaufeld et al., 2020; Martin, 2020; ten Oever et al., 2022), and in line with the idea that the timescale of syntactic processing overlaps with the delta frequency range (Henke & Meyer, 2021; Meyer et al., 2020; see also Figure 3), we focus on MEG activity in this band.

Previous work has identified several brain areas in the left hemisphere that are responsive to complexity metrics derived from incremental parse steps, including the inferior frontal lobe (Bhattasali et al., 2019; Giglio et al., 2024; Nelson et al., 2017), the anterior temporal lobe (Bhattasali et al., 2019; Brennan & Pylkkänen, 2017; Brennan et al., 2012, 2016; Nelson et al., 2017), and posterior superior and middle temporal areas (Brennan et al., 2016; Giglio et al., 2024; Li & Hale, 2019; Nelson et al., 2017; Lopopolo et al., 2021). Assuming that these effects reflect the operations involved in syntactic structure building and that people employ similar parsing strategies when processing English and Dutch sentences, we expect effects in the same brain regions. Because the majority of this work has used fMRI, the timing of these effects is less clear, though the results of electrophysiological studies suggest that effects of structure building are reflected in brain activity within the first 500 ms after word onset (Brennan & Pylkkänen, 2017; Hale et al., 2018). In sum, the predictive accuracy of the different parsing models can give us insight into the (potentially language-specific) processing strategies used by people comprehending Dutch (i.e., *how* they build structure), and the spatial-temporal properties of the effects provide clues about *when* and *where* in the brain these processes are implemented.

## 2. Methods

### 2.1 Participants

Twenty-four right-handed native speakers of Dutch (18 female, mean age = 33.4 years, age range = 20-58 years) were recruited via the SONA system of Radboud University Nijmegen. They all reported normal hearing, had normal or corrected-to-normal vision, and did not have a history of language-related impairments. Participants gave written informed consent to take part in the experiment, which was approved by the Ethics Committee for human research Arnhem/Nijmegen (project number CMO2014/288).

### 2.2 Stimuli

The stimuli consisted of stories from three fairy tales in Dutch: one story by Hans Christian Andersen and two by the Brothers Grimm. All stories contain a rich variety of naturally occurring sentence structures, varying in syntactic complexity. In total, there are 8551 words in 791 sentences, which are on average 10.8 words long (range = 1-35, SD = 5.95). They were auditorily presented in nine separate segments, all of which were between 289 (4 min, 49 sec) and 400 seconds long (6 min, 40 sec; see Supplementary Information S1), for a total of 49 minutes and 17 seconds. The loudness of each audio segment was normalized using the Ffmpeg software (EBU R128 standard). The transcripts of each story were automatically aligned with their corresponding audio recording using the WebMAUS segmentation software (Kisler, Reichel, & Schiel, 2017).

### 2.3 Syntactic annotations

We manually annotated syntactic structures for all sentences in the audiobooks following an adapted version of X-bar theory (Carnie, 2021; Jackendoff, 1977). To be specific, we consistently created an X-bar structure for NPs and VPs, whereas intermediate projections for all other phrases were drawn only if they were needed to attach adjacent words to the structure (e.g., APs were unbranched unless they were modified by an adverb or prepositional phrase). The X-bar template for NPs and VPs was strictly enforced in order to make a structural distinction between arguments and adjuncts; arguments were attached as sister of the head, while adjuncts were attached to an intermediate projection (Jackendoff, 1977). All phrases except for VPs and IPs are head-initial. An example of a hierarchical structure is given in Figure 2.

**Figure 2.**
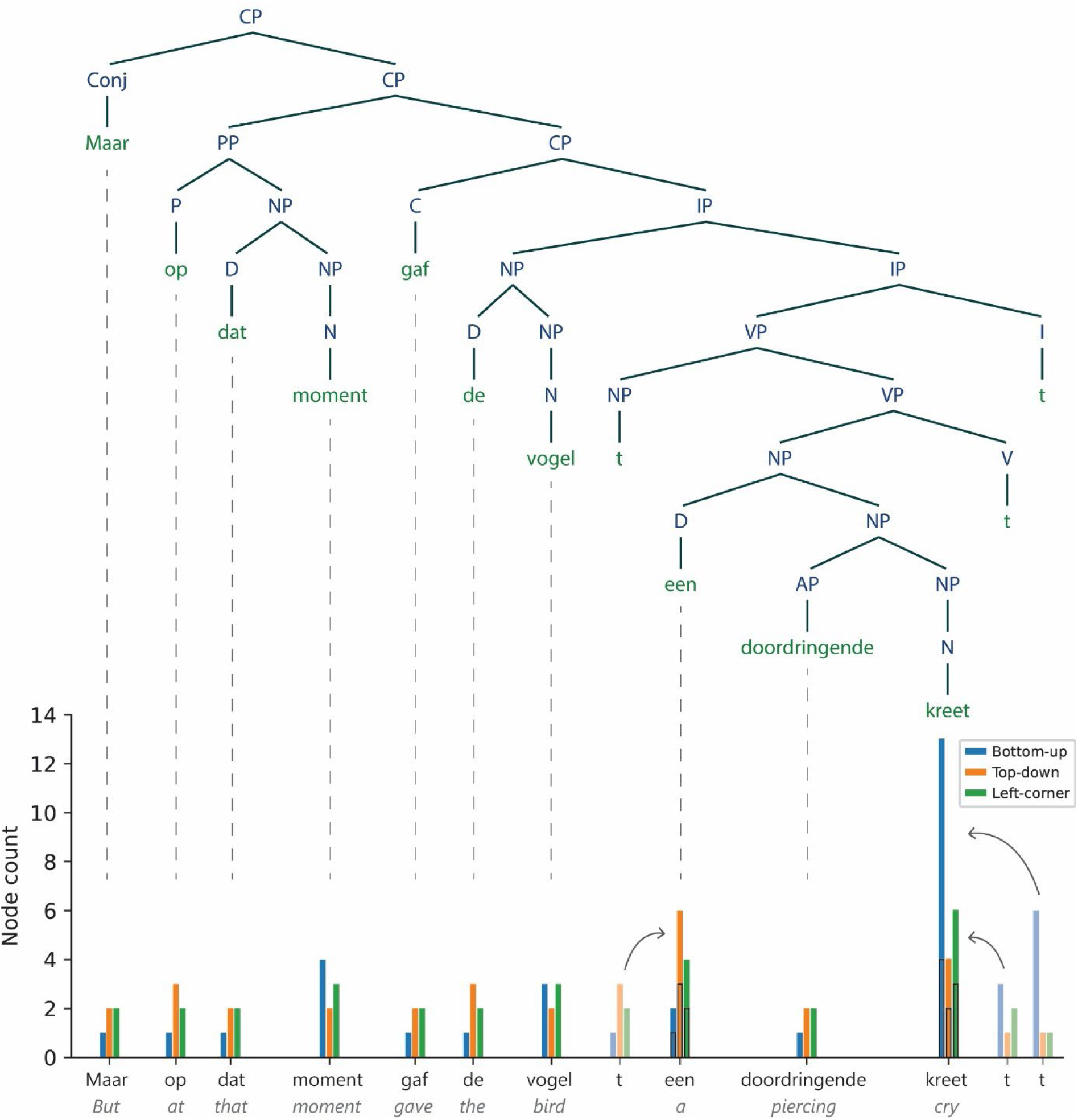
Syntactic structure of an example sentence from one of the audiobook stories, with node counts for each terminal node presented below. The *t* stands for trace and refers to the position at which the word or phrase to which it is related is interpreted. It does not have an acoustic correlate in the speech signal. Node counts for traces are added to node counts of the next word, except for traces that occupy sentence-final position, whose node counts are assigned to the previous word.

Node counts were computed for each word in every sentence in three different ways. On the bottom-up strategy, a constituent node is posited when all daughter nodes have been encountered. This amounts to counting the number of closing brackets directly following a given word in a bracket notation. On the top-down strategy, a node is posited right before it is needed to attach the upcoming word to the structure. This amounts to counting the number of opening brackets directly preceding a given word in a bracket notation. And on the left-corner strategy, a node is posited when its left-most daughter node has been encountered. Terminal nodes were not included in the node count calculation.

As can be seen in Figure 2, the X-bar structures contain traces of movement (Carnie, 2021). As these empty elements do not have an acoustic correlate in the speech signal, we assigned their node counts to another word in the same sentence. Our reasoning was that the precise location of these elements could not be predicted with certainty, though it could be inferred after their putative position. We therefore decided to add the node count of each empty element to the node count of the word following it. By doing so, we aimed to capture the syntactic processes associated with these elements (e.g., long-distance dependency resolution) around the times they occur. We will come back to this point in the discussion. The resulting node counts were time-aligned with the onsets of the words in the audio recordings. Each audio file could thus be represented as a vector with a node-count value at the onset of each word and zeros everywhere else. The vectors for each of the three parsing strategies were the predictors in the TRF analysis (see Section 2.7).

It has been suggested that the timescale of syntactic processing overlaps with the delta frequency range (Henke & Meyer, 2021; Meyer et al., 2020), which is consistent with the syntactic information rate in the nine story segments that were used in this study. Syntactic information rates were derived from the time between the onsets of consecutive words with node counts above 2. Because we aim to illustrate the rate at which syntactic structure building can be initiated in the stimuli, only syntactically demanding words (i.e., node count > 2) were included in the rate calculation. The probability density plots in Figure 3 show that the syntactic information rate in all story segments largely falls within the delta range, supporting the idea that this is the timescale at which syntax can be perceptually inferred from naturalistic speech.

**Figure 3.**
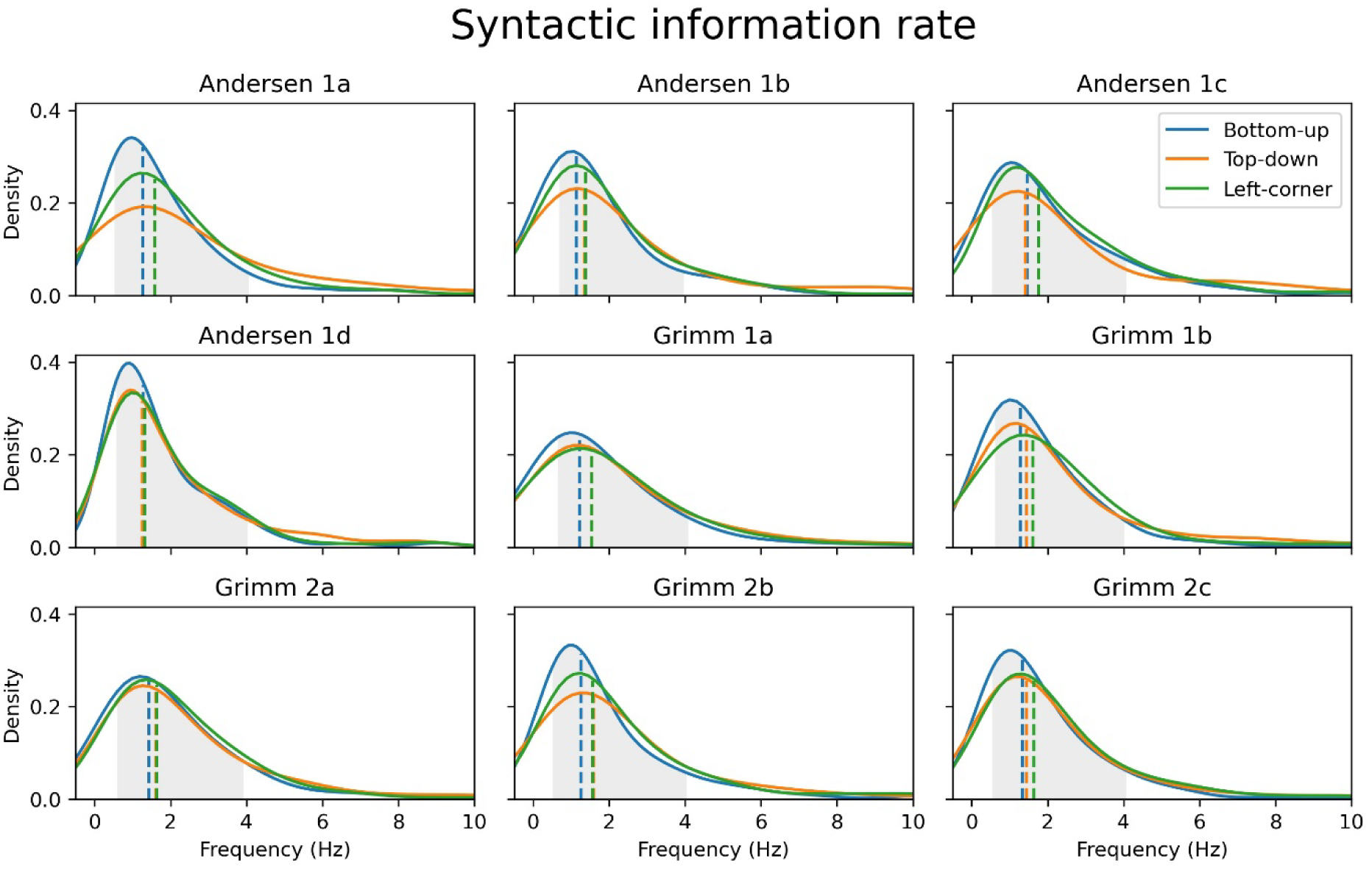
Probability density functions showing that the average rate at which syntactically relevant information is presented largely falls within the 0.5-4 Hz delta band (highlighted in grey) in all story segments. The dashed vertical lines reflect the median rate per parsing method.

### 2.4 Procedure and data acquisition

Participants were individually tested in a magnetically shielded room. They were instructed to attentively listen to the nine audiobook stories while sitting still and looking at a fixation cross that was presented in the middle of the screen. After each of the nine story blocks, five multiple-choice comprehension questions (each with four options) were asked. On average, participants answered 88.1% of the questions correctly (SD = 7.52%), showing that they paid attention to and understood the content of the stories.

The MEG data were recorded with a 275-channel axial gradiometer CTF system at a sampling rate of 1200 Hz. The audio recordings were presented using Psychtoolbox in MATLAB (Brainard, 1997) via earphones inserted into the ear canal. Participants’ eye movements and heartbeat were recorded with EOG and ECG electrodes, respectively. Throughout the recording session, their head position was monitored using three head localization coils, one placed in fitted earmolds in each ear and one placed at the nasion (Stolk et al., 2013). Each block started with a 10-second period during which resting state data were recorded. In the break between story blocks, participants were instructed to reposition their head location in order to correct for head movements. After the MEG session, each participant’s head shape was digitized using a Polhemus 3D tracking device, and their T1-weighted anatomical MRI was acquired using a 3T Skyra system (Siemens).

### 2.5 MEG preprocessing

Preprocessing was done using MNE-Python (version 0.23.1). The MEG data were first down-sampled to 600 Hz and band-pass filtered at 0.5-40 Hz using a zero-phase FIR filter (MNE-Python default settings). We then interpolated channels that were considered bad using Maxwell filtering, and used Independent Component Analysis to filter artifacts resulting from eye movements (EOG) and heartbeats (ECG). We segmented the data into nine large epochs, whose onsets and offsets corresponded to those of the audio recordings. Source reconstruction was done for each epoch separately.

### 2.6 Source reconstruction

Individual head models were created for each participant with their structural MRI images using FreeSurfer (Dale et al., 1999). The MRI data were then co-registered to the MEG with MNE co-registration, using the head localization coils and the digitized head shape. We set up a bilateral surface-based source space for each individual participant using fourfold icosahedral subdivision, resulting in 2562 continuous source estimates in each hemisphere. The forward solution was computed using a BEM model with single layer conductivity. We low-pass filtered the signal at 4 Hz using a zero-phase FIR filter (corresponding to the 0.5-4 Hz delta band) and estimated sources using the dSPM method (noise-normalized minimum norm estimate), with source dipoles oriented perpendicularly to the cortical surface. The noise covariance matrix was calculated based on the resting state data that were recorded before each story (all concatenated). Before TRF analysis, each source estimate was downsampled to 100 Hz to speed up further computations.

### 2.7 Predictor variables

To control for brain responses to acoustic information, all models included two acoustic predictors: an eight-band gammatone spectrogram (i.e., envelope of the acoustic signal in different frequency bands) and an eight-band acoustic onset spectrogram. Both spectrograms covered frequencies from 20 to 5000 Hz in equivalent rectangular bandwidth space (Heeris, 2018), and were resampled to 100 Hz to match the sampling rate of the MEG data. The onset spectrogram was derived from the gammatone spectrogram using an auditory edge detection model (Brodbeck et al., 2020) implemented in Eelbrain (Version 0.37.3; Brodbeck et al., 2021).

All models also included four word-based predictors that are all strongly linked to brain activity during naturalistic language processing (Brodbeck et al., 2018, 2022; Weissbart et al., 2020). These were word rate and the three statistical predictors word frequency, surprisal, and entropy. All predictors were modeled on the gammatone predictors in terms of length and sampling rate.

The word rate predictor is simply a one-dimensional array with the value 1 at word onsets and the value 0 everywhere else.

The frequency of word *w* was computed by the taking negative logarithm of the number of occurrences of *w* of per million words, extracted from the SUBTLEX-NL database of Dutch word frequencies (Keuleers et al., 2010):

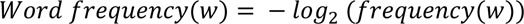

Word frequency was represented via the negative logarithm, because in this way infrequent words will get high values and frequent words will get low values, in line with the brain response to word frequency (i.e., a larger response to infrequent words; Brennan et al., 2016; Brodbeck et al., 2018). For some words we could not compute a frequency value because the word did not appear in the database. Manually checking them revealed that these were uncommon (and thus likely infrequent) words, so we assigned to them the value corresponding to the lowest frequency of all words present in the audiobook.

Surprisal is the conditional probability of a word given the preceding linguistic context, quantified as the negative log of this probability (Hale, 2001, 2014). Thus, the surprisal of word *w* at position *t* is calculated via:

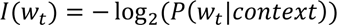

Word surprisal was computed from conditional probabilities obtained with GPT-2 for Dutch (De Vries & Nissim, 2021). GPT-2 used the preceding 30 words as context, so *context* in the formula above refers to (𝑤_𝑡−30_ … 𝑤_𝑡−1_).

Entropy at word position *t* is the uncertainty before observing the next word *w*_t+1_ given the preceding context. Context was again defined as the previous 30 words (including *w*_t_), and conditional probabilities were again obtained with GPT-2 for Dutch (De Vries & Nissim, 2021). Entropy at word position *t* was then calculated as the sum of the conditional probabilities of each next word (within the set of possible upcoming words *W*), weighted by the negative logarithm of this probability:

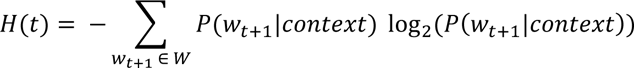

Our syntactic models included the syntactic predictors bottom-up node count, top-down node count and left-corner node count. We constructed a total of four models (see Table 2).

**Table 2.**
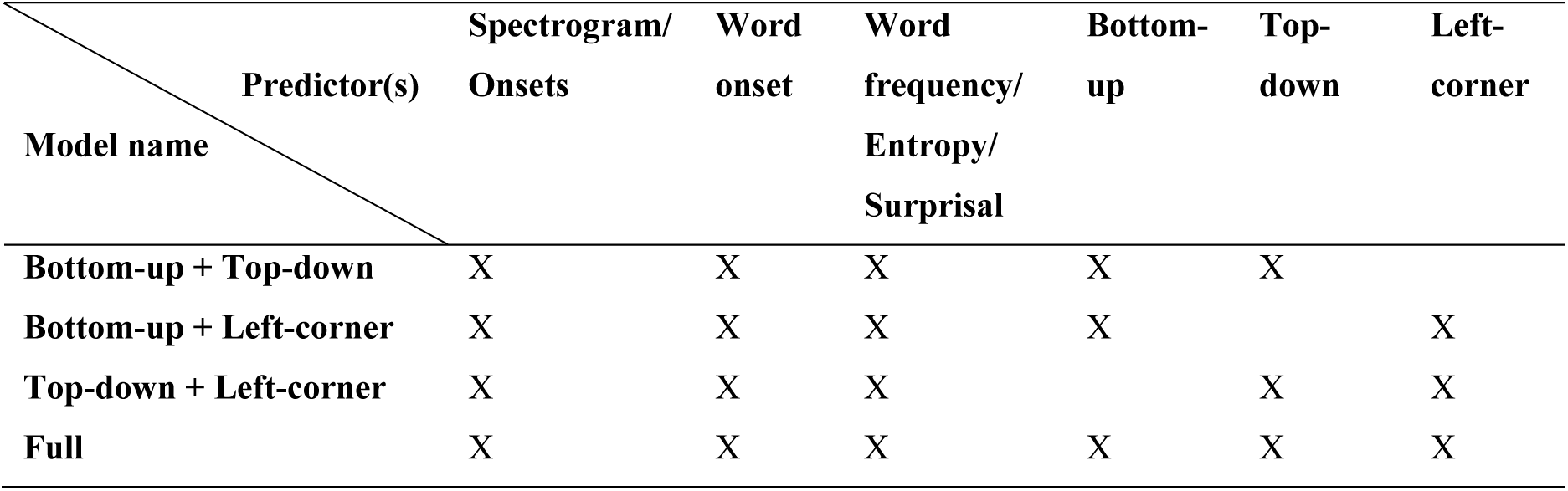
Predictors included in each model.

| Predictor(s) | Spectrogram/<br>Onsets | Word<br>onset | Word<br>frequency/<br>Entropy/<br>Surprisal | Bottom-up | Top-down | Left-corner |
| --- | --- | --- | --- | --- | --- | --- |
| **Bottom-up + Top-down** | X | X | X | X | X |  |
| **Bottom-up + Left-corner** | X | X | X | X |  | X |
| **Top-down + Left-corner** | X | X | X |  | X | X |
| **Full** | X | X | X | X | X | X |

### 2.8 Model estimation

TRFs were estimated for each subject and MEG source point separately using Eelbrain (Version 0.37.3; Brodbeck et al., 2021). The MEG response at time *t*, denoted as *ŷ*(*t*), was predicted jointly by convolving each TRF with a predictor time series shifted by K time delays (Brodbeck et al., 2021, 2022):

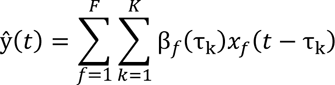

Here, *x*_f_ is the predictor time series and β*_f_*(τ_k_) is the coefficient of the TRF of the corresponding predictor at delay τ_k_. The coefficient of the TRF at delay τ thus indicates how a change in the predictor affects the predicted MEG response τ milliseconds later. To generate each TRF, we used 50-ms wide Hamming windows and shifted the predictor time series between −100 and 1000 ms at a sampling rate of 100 Hz, thus yielding K = 110 different delays. The length of the TRF was chosen based on the latency of syntactic effects in naturalistic paradigms (Brennan & Hale, 2019; Hale et al., 2018). Before estimating the TRF, all predictors as well as the MEG data were mean-centered and then normalized by dividing by the mean absolute value.

TRFs were estimated using a five-fold cross-validation procedure. We first concatenated the data for each subject along the time axis, and then split them up into five equally long segments. During each cross-validation run, three segments were used for training, one for validation, and one for testing. For each test segment, there were four training runs with each of the remaining segments serving as the validation segment once. Using a boosting algorithm (David et al., 2007) to minimize the l1 error, one TRF was estimated for each of the four training runs (selective stopping based on l1 error increase). The resulting four TRFs were averaged to predict responses in the test segment. This analysis yields an average TRF for each predictor in each model, as well as a measure of reconstruction accuracy for the whole model. Reconstruction (or predictive) accuracy refers to the fit between the predicted and the observed MEG signal at each source point, quantified in terms of explained variance in R^2^. Reconstruction accuracy can be seen as a measure of neural tracking: the larger the reconstruction accuracy for a given model, the more closely the brain tracks the predictors in that model.

### 2.9 Model comparison

We first tested whether separately adding each of the three syntactic predictors to a base model with all control predictors (see Table S2.1 in the Supplementary Information) would increase the reconstruction accuracy. Having established this (see Supplementary Information S2.1), we then determined the unique contribution of each syntactic predictor by comparing the reconstruction accuracy of the full model to the reconstruction accuracy of a null model from which only one of the predictors was omitted. The three null models we evaluated were Bottom-up + Top-down, Bottom-up + Left-corner, and Top-down+ Left-corner (see Table 2). Comparing their reconstruction accuracy to the accuracy of the full model yields an accuracy difference measure for the left-corner, top-down, and bottom-up predictors, respectively. This comparison thus tests whether a predictor explains variance in the brain signal above and beyond the variance explained by all other predictors.

To determine where in the brain the reconstruction accuracy of the full model was different from that of the null model, we smoothed the source points of both models separately (Gaussian window, SD = 14 mm) and tested for differences in their source maps using non-parametric cluster-based permutation tests (Maris & Oostenveld, 2007). For all contrasts between full and null model, we applied two-tailed paired-samples t-tests at each source point, clustered adjacent source points (minsource = 10) with an uncorrected p-value lower than 0.05, and evaluated clusters of activity by comparing their cluster-level test statistic (sum of individual t-values) to a permutation distribution. The permutation distribution was generated based on the maximum cluster-level t-value in each of 10,000 random permutations of the same data, in which the condition labels were shuffled within subjects. The significance of clusters was evaluated at an alpha value that was Bonferroni-corrected for the number of tests (alpha = 0.05/n_tests_). As an estimation of the effect size of the significant clusters, we report t_av_, which corresponds to the average t-value within the significant cluster (the cluster-level t-value divided by the number of significant source points). When multiple clusters are significant, we report the test statistic of the cluster with the largest number of significant source points.

The syntactic predictors were quite highly correlated (r_BU-TD_ = 0.33, r_LC-BU_ = 0.74, r_TD-LC_ = 0.80; see Supplementary Information S2), potentially leading to multicollinearity, which affects estimation of the TRF coefficients. To control for this possibility, we separately compared the Bottom-up, Top-down, and Left-corner models to a base model, which included no syntactic predictors at all (see Table S2.1 in the Supplementary Information). This analysis does not suffer from multicollinearity issues because the correlated predictors never appear in the same model. The results of this analysis are both qualitatively and quantitively very similar to those reported in the main analyses (see Supplementary Information S2.1-S2.3), supporting the conclusion that the different predictors explain unique variance in the MEG data despite being positively correlated.

### 2.10 Evaluation of response functions

In addition to analyzing the fit between the predicted and the observed signal, we also evaluated the estimated response function, which provides information about the temporal relationship between the predictor and the neural response. This analysis involves the coefficients of the TRF at each time and source point (sources smoothed by a Gaussian window, SD = 14 mm). If the coefficients for a given predictor are significantly non-zero, this indicates that the brain responds to the information encoded in that predictor. In a spatiotemporal cluster-based permutation analysis, we first applied two-tailed one-sample t-tests at each source-time point to determine whether the TRF coefficients deviate from zero. The t-values of adjacent source-time points (minsource = 10, mintime = 40 ms) with an uncorrected p-value lower than 0.05 were then summed, and their cluster-level test statistic was compared to a permutation distribution based on 10,000 random permutations of the same data. The significance of clusters was evaluated at an alpha value (of 0.05) that was Bonferroni-corrected for the number of tests.

## 3. Results

### 3.1 Model comparison

Using cluster-based permutation tests in source space, we tested where in the brain the reconstruction accuracies were modulated by each of the three syntactic predictors. Clusters of activity were evaluated at alpha = 0.0083 (n_tests_ = 3 accuracy differences * 2 hemispheres). All predictors significantly improve reconstruction accuracy, with clusters mostly in the left hemisphere (see Figures 4A-C). The improvement is largest for the top-down predictor, which explains variance in many regions of the left hemisphere (t_av_ = 5.49, p = .0034), as well as in an anterior part of the right temporal lobe (t_av_ = 2.60, p = .0075). The strongest left-hemispheric effects are found in superior and middle temporal regions and in inferior and middle frontal regions. The bottom-up predictor similarly engages inferior frontal and temporal regions, but only one cluster around Heschl’s gyrus is significant at the adjusted alpha level (t_av_ = 3.84, p = .0081). Last, the left-corner predictor improves reconstruction accuracy in an area at the border of the temporal and frontal lobe in both the left (t_av_ = 5.72, p = .0057) and the right hemisphere (t_av_ = 3.35, p = .0081). It is noteworthy that even though the three syntactic predictors are positively correlated with one another, each of them explains unique variability in the MEG data that could not be attributed to the other two syntactic predictors.

**Figure 4.**
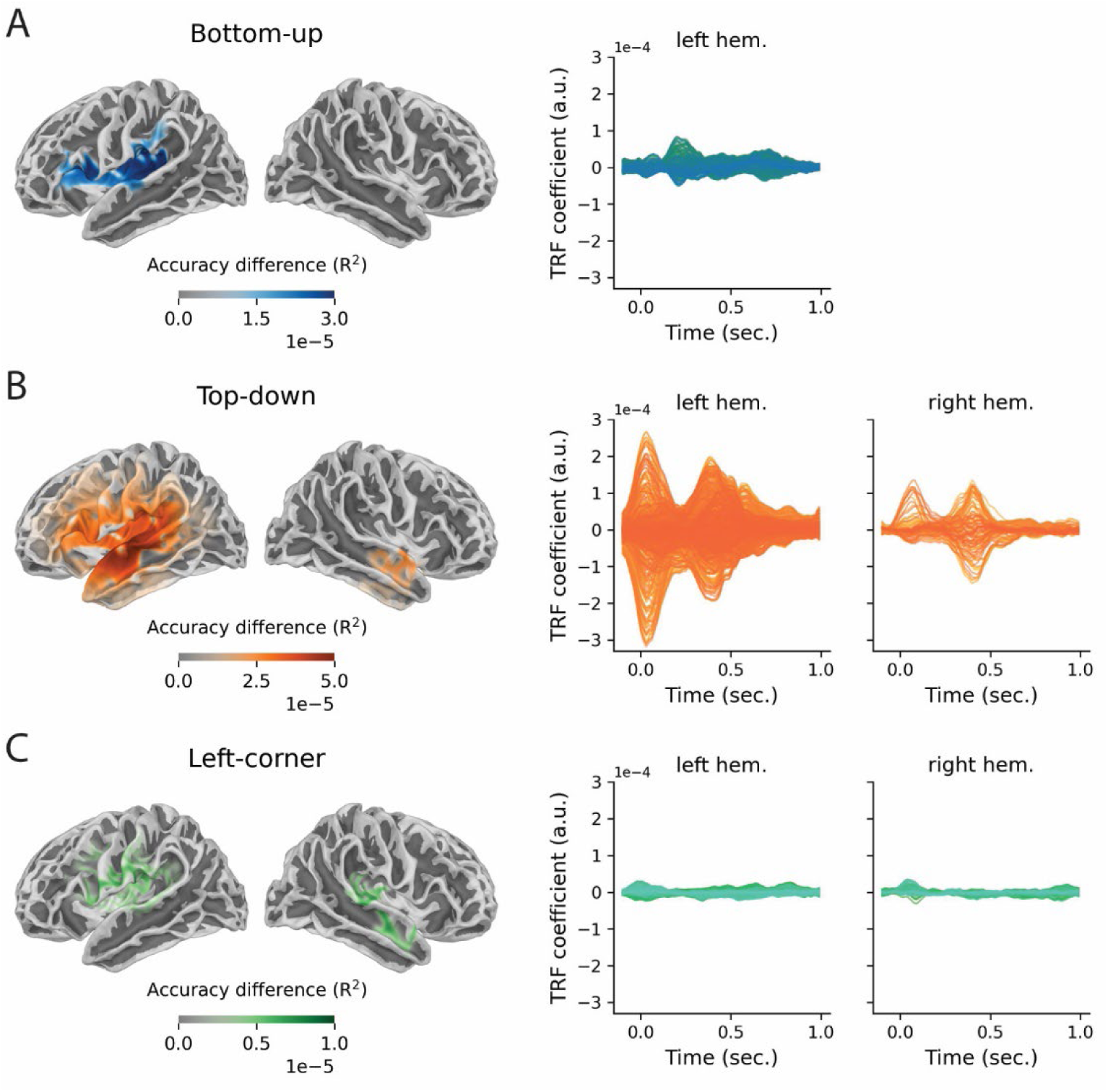
Sources of significantly improved explained variance and temporal response functions in significant source points. Results are shown separately for the effects of the bottom-up **(A)**, top-down **(B)** and left-corner **(C)** predictors. Significance was determined by comparing the reconstruction accuracy of the full model to the reconstruction accuracy of a null model from which the relevant predictor was omitted. All clusters that were significant at uncorrected alpha = 0.05 are displayed. Notice that the scales of the color bars are different across the source plots.

### 3.2 Evaluation of the response functions

Given that all of the predictors increase reconstruction accuracy, we examined their response functions, which reveal a more detailed picture of the time course of the neural response to syntactic information. Figure 4 shows the TRFs within the significant regions of the cluster-based source analysis of reconstruction accuracies, in the left and the right hemisphere separately. These plots yield an estimate of the magnitude of the brain response to the syntactic predictor at each time point. All TRFs come from the full model, in which all predictors are competing for explaining variance, so the increases in amplitude reflect components of the neural response that are best explained by the respective predictor. In line with the reconstruction accuracy results, the TRF amplitudes clearly reveal that the neural response to the information encoded in top-down node counts is stronger than the response to node counts derived from the bottom-up or the left-corner method.

While these results are informative about the overall strength and timing of the responses, they do not show which regions are involved at which time points. This information is provided in Figure 5, which shows the source t-values (based on two-tailed, one-sample t-tests) of the TRFs for the three syntactic predictors, split up into four time windows (corresponding to different delays in TRF estimation). Non-parametric cluster-based permutation tests were used to determine when and where the TRF coefficients of each syntactic predictor deviated from zero. Because this involves 6 comparisons (3 TRFs * 2 hemispheres), clusters were evaluated at alpha = 0.0083. Focusing on left-hemispheric sources, this analysis revealed a negative cluster for the top-down predictor, broadly distributed in frontal and temporal regions (t_av_ = −0.53, p < .001)^2^, and a positive cluster around the parahippocampal gyrus and anterior temporal lobe (t_av_ = 0.68, p < .001). The strongest effect of the bottom-up TRF was in a region centered around the left inferior frontal cortex (t_av_ = 0.43, p < .001), and the TRF of the left-corner predictor peaked in (anterior) temporal (t_av_ = −0.38, p = .0062) and frontal regions (t_av_ = 0.48, p < .001). In sum, these results show the dynamic nature of structure building: the relationship between the predictor and certain brain regions is time-dependent, suggesting that these areas are differently involved across different delays.

**Figure 5.**
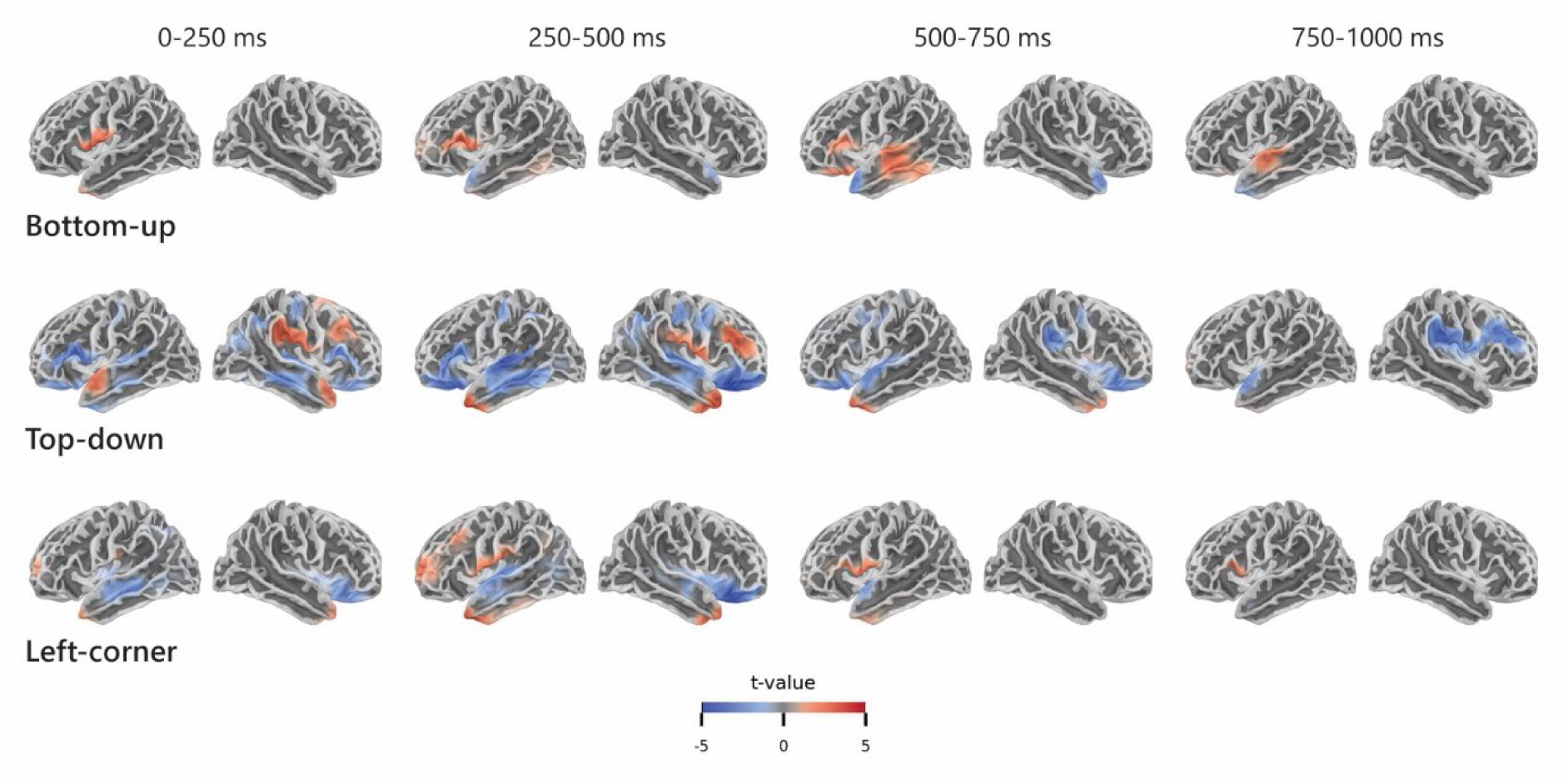
Sources of the TRFs for node count derived from bottom-up, top-down, and left-corner parsers, representing early to late responses. The colors represent positive (red) or negative (blue) t-values in sources that were significantly responsive (at corrected alpha = 0.0083) to the predictor in the indicated time windows.

### 3.3 Region of interest analysis

To further explore the spatiotemporal differences between the response functions of the syntactic predictors, we analyzed the TRFs in three specific regions of interest (ROIs) that have been linked to syntactic structure building in naturalistic contexts. These ROIs are the inferior frontal gyrus (IFG), posterior temporal lobe (PTL) and anterior temporal lobe (ATL) in the left hemisphere (Figure 6A), all of which also showed up in one or more of the contrasts in the accuracy analysis (Figure 4) and the TRF analysis (Figure 5). The locations of these ROIs were defined based on the peak coordinates in Montreal Neurological Institute (MNI) space of an fMRI study on syntactic structure building (Pallier et al., 2011). We used these coordinates as seeds to create spheres with a 40-mm radius for the temporal pole (MNI coordinates: −48, 15, −27) and posterior superior temporal sulcus (−51, −39, 3). In addition, 30-mm spheres were created around the coordinates of the left inferior frontal cortex, which included both the pars triangularis (−51, 21, 21) and the pars orbitalis (−45, 33, −6).

**Figure 6.**
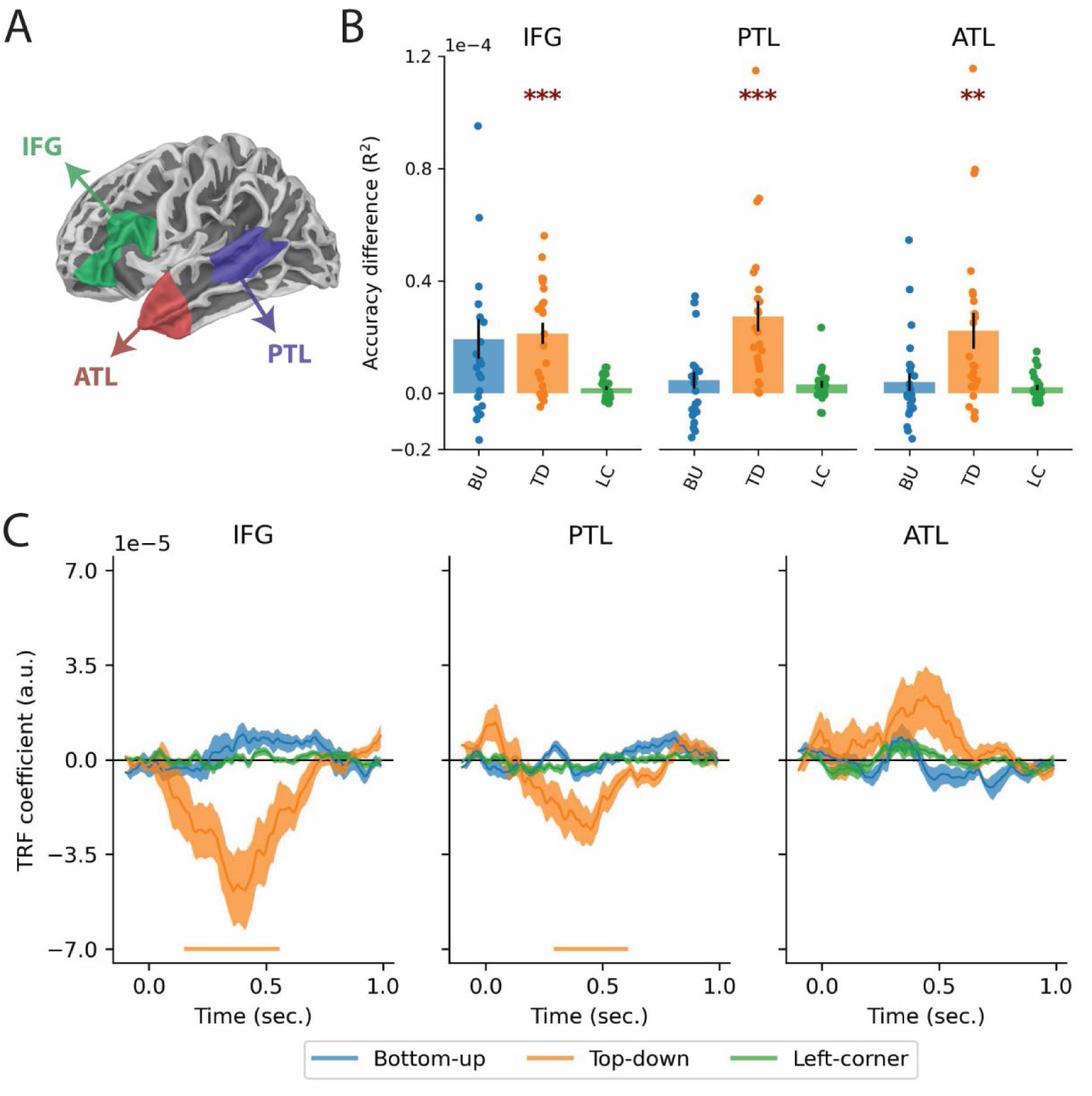
Region of interest analysis. **(A)** Spatial extensions of the three regions of interest. **(B)** Difference in reconstruction accuracy with the full model, plotted for left IFG, PTL, and ATL. The labels on the x-axis refer to the syntactic predictors that were taken out of the full model, so the height of each bar indicates the reduction in reconstruction accuracy compared to the full model when only that predictor is omitted. The drops represent the accuracy difference for individual participants, and the error bars represent the standard error of the mean across subjects. **(C)** Temporal response functions for node count derived from bottom-up, top-down, and left-corner parsers in the full model. Error bars reflect the standard error of the mean per time sample. The horizontal bars below the TRFs reflect the time points at which the TRFs were significantly non-zero.

The results in Figure 6B show for each ROI the improvement in reconstruction accuracy when the syntactic predictors are separately added to the null model. Effects of the top-down predictor are strongest in general, and in particular in the PTL, where all subjects show evidence of responses associated with top-down node counts. Supporting this impression, two-tailed paired-samples t-tests (with alpha = 0.0056, Bonferroni-corrected for 9 tests) reveal that adding the top-down predictor to the null model improves the reconstruction accuracy in all three ROIs (IFG: t(23) = 5.54, p < .001; PTL: t(23) = 5.04, p < .001; ATL: t(23) = 3.50, p = 0.0020). The addition of the bottom-up predictor seems to improve reconstruction accuracy in the IFG, but this effect did not survive multiple-comparisons correction, t(23) = 2.73; p = 0.012.

In each of the three ROIs, we used cluster-based permutation tests to determine when the TRFs of each syntactic predictor deviated from zero. Again, this involves 9 comparisons (3 TRFs * 3 ROIs), so clusters were evaluated at alpha = 0.0056. As shown in Figure 6C, the TRF of the top-down predictor showed negative-going modulations in the IFG (from 160 to 560 ms, t_av_ = −2.91, p < .001) and PTL (from 300 to 610 ms, t_av_ = −3.04, p < .001). As in the reconstruction accuracy analysis above, the modulation of the bottom-up TRF in the IFG (from 590 to 730 ms, t_av_ = 2.61, p = .015) was not significant at the adjusted alpha level. It is noteworthy that the TRFs within the same region are not consistent across different predictors. While their sign is not directly interpretable, the differences in Figure 6C (e.g., the consistently positive TRF for bottom-up vs. the negative TRF for top-down in the IFG) indicate that the processes these regions are involved in are not the same for the different syntactic predictors.

## 4. Discussion

In this study, we investigated when the brain projects its knowledge of syntax onto speech during natural story listening. Atemporal syntactic structures were expressed in the time domain via the use of incremental node count, whose temporal distribution showed that the delta band is a syntactically relevant timescale in our stimuli. Using a forward modelling approach to map node counts onto source-reconstructed delta-band activity, we then compared three parsing models that differ in the dynamics of structure building. A key finding of these analyses is that neural source dynamics most strongly reflect node counts derived from a top-down parsing model. This model postulates syntactic structure early in time, suggesting that predictive structure building is an important component of Dutch sentence comprehension. The additional (though weaker) effects of the bottom-up and left-corner predictors indicate that integratory and mildly-predictive parsing also play a role, and suggest that people’s processing strategy might be flexibly adapted to the specific properties of the linguistic input.

### 4.1 Predictive structure building in the brain

Node counts derived from all three parsing models explained unique variance in delta-band MEG activity, consistent with recent studies showing a relationship between delta-band activity and syntactic processing (Bai et al., 2022; Brennan & Martin, 2020; Coopmans et al., 2022; Ding et al., 2016; Henke & Meyer, 2021; Kaufeld et al., 2020; ten Oever et al., 2022). The results also align with prior studies that use neuro-computational language models to study syntactic processing (Bhattasali et al., 2019; Giglio et al., 2024; Nelson et al., 2017 ; Brennan & Pylkkänen, 2017; Brennan et al., 2012, 2016; Li & Hale, 2019; Lopopolo et al., 2021), but they nevertheless advance our understanding of the neural encoding of syntax due to the inclusion of low-level (linguistic and non-linguistic) predictors in our TRF models. In addition to a number of acoustic predictors, the null models also contained information-theoretic predictors (e.g., surprisal, entropy) that partially reflect syntactic information and therefore explain some of the variance in brain activity whose origin is syntactic. By including these semi-syntactic control predictors, we stacked the cards against us and likely underestimated the ‘true’ neural response to syntactic structure. The fact that we nevertheless find robust effects of syntactic processing underscores the relevance of syntax for the neural mechanisms underlying language comprehension.

Of all syntactic predictors, node counts derived from a top-down parsing method were the strongest syntactic predictor of brain activity in language-relevant areas. These effects peaked twice within the first 500 ms after word onset and encompassed mostly inferior and middle frontal, and superior and middle temporal areas in the left hemisphere. The predictiveness of top-down node counts is somewhat at odds with previous studies that have looked at different parsing models in naturalistic comprehension, which either find that top-down methods are less predictive of brain activity (Giglio et al., 2024; Nelson et al., 2017; Slaats et al., 2024) or that they do not differ from other parsing methods (Brennan et al., 2016). What could account for the strong top-down effects? One explanation is that top-down node counts capture the predictive nature of language processing well. There is substantial evidence from psycholinguistics that people generate structural predictions across a variety of syntactic constructions (Arai & Keller, 2013; Ferreira & Qiu, 2021; Lau et al., 2006; Staub & Clifton, 2006; Yoshida et al., 2013), and they do so in naturalistic contexts as well (Brennan & Hale, 2019; Hale et al., 2018; Heilbron et al., 2022). These predictive structure-building processes are mostly associated with activity in the left posterior temporal lobe (PTL; Brennan et al., 2016; Matchin et al., 2017, 2019; Matar et al., 2021; Nelson et al., 2019) and left inferior frontal gyrus (IFG; Matchin et al., 2017), both of which were responsive to the top-down predictor in the current study. Matchin et al. (2019) suggest that the PTL is involved in predictive activation of sentence-level syntactic representations and/or increased maintenance of the syntactic representations associated with lexical items when they are presented in a sentential context (see also Matar et al., 2021). On both interpretations, the PTL encodes structural representations that can be activated in a predictive fashion and are later to be integrated with the sentence-level syntactic representation in IFG (Hagoort, 2005; Hagoort & Indefrey, 2014; Snijders et al., 2009). Importantly, this process does not proceed in a purely feedforward manner, but rather relies on recurrent connections between temporal and frontal regions (Hultén et al., 2019). That the TRF for the top-down predictor is bimodal is consistent with this idea (Figure 4B). The first peak, observable in the PTL only (Figure 6C), likely reflects predictive structure building, which can occur in the PTL within the first 150 ms after word onset (Matchin et al., 2017). Indeed, the early timing of this effect is consistent with the predictive nature and temporal spell-out of top-down parsing, which is maximally eager and postulates syntax early in time. The second, later peak is largely consistent with syntactic surprisal effects in recent naturalistic M/EEG studies (Brennan & Hale, 2019; Hale et al., 2018; Heilbron et al., 2022), and might reflect responses related to the (dis)confirmation of predicted structures based on incoming information. An additional observation is that the effects of the top-down predictor, and to a lesser extent also those of the left-corner predictor, are somewhat bilateral. A tentative explanation for this finding is that syntactic prediction, as quantified by top-down and left-corner node counts, is demanding and therefore requires support from right-hemispheric regions. These areas are presumably not the locus of the syntactic representations and computations themselves but might be activated when processing demands are increased (Hartwigsen, 2018).

The bottom-up TRF was quite specifically modulated in the left inferior frontal cortex (Figure 5), in line with previous studies that found this area to be responsive to parametric manipulations of syntactic structure (Giglio et al., 2022; Matchin et al., 2019; Pallier et al., 2011; ten Oever et al., 2022; Zaccarella et al., 2017). Compared to the top-down TRF, however, bottom-up effects were relatively weak in amplitude. One possibility is that the bottom-up and top-down effects are inversely related, such that strong top-down effects, which indicate predictive processing, are accompanied by weak bottom-up effects, and vice versa. One of the advantages of prediction in language comprehension is that it can reduce the burden on future integration processes (Kuperberg & Jaeger, 2016). If a structural representation has already been pre-built or pre-activated, correctly predicted incoming words only have to be inserted into the existing structure, so integration costs for these words are low. Because integration costs are approximated via bottom-up node count, any effects of bottom-up node count should be reduced if people successfully engage in predictive processing. This account would thus predict that when top-down metrics modulate brain activity for a given sentence, bottom-up metrics will not provide a good fit for that same sentence, and conversely, when top-down metrics do not provide a good fit, bottom-up metrics should be highly predictive. While it should be investigated in future work whether the effects of top-down and bottom-up metrics indeed go hand in hand in this anti-correlated way, the results of two naturalistic studies with spontaneous speech are consistent with this possibility. Contrasting with coherent audiobook narratives, spontaneously produced speech contains dysfluencies and corrections, which might make participants less inclined to rely on predictive processing. In an fMRI study by Giglio et al. (2024), English speakers had to listen other people’s verbal summaries of a tv episode. The authors found that bottom-up node counts modulated activity in language-relevant brain areas (LIFG and LPTL) more strongly than top-down node counts. Second, an EEG study by Agmon et al. (2023) investigated the neural encoding of different linguistic features when Hebrew speakers listened to a spontaneously generated narrative. Using a TRF analysis on low-frequency activity, they found that the neural response for words closing a clause (bottom-up) was stronger than the neural response for words opening a clause (top-down). Both studies indicate that in the comprehension of spontaneous speech the brain relies relatively strongly on integratory processing. A possible implication of these findings is that parsing strategies can be flexibly adapted to the specific properties of the current linguistic input (e.g., grammatical properties, sentence complexity, reliability of predictive cues), such that people are less likely to engage in predictive structure building if the input contains ungrammatical sentences that make predicting ineffective (Brothers et al., 2019; Kuperberg & Jaeger, 2016). As an exploratory analysis of the potential trade-off between predictive and integratory structure building, we investigated whether predictability (quantified through surprisal) modulates the demands on bottom-up structure building (see the Supplementary Information S3; see also Slaats et al., 2024). How exactly the demands on predictive and integratory structure building dynamically change over the course of sentence processing should be investigated in future research.

Effects of the left-corner predictor were also relatively weak compared to effects of the top-down predictor, in terms of both reconstruction accuracy and TRF amplitude. The comparatively weak effects of left-corner node counts might indicate that left-corner metrics are insufficiently predictive to account for the comprehension of head-final constructions of Dutch. The left corner of head-initial structures is very informative, which could explain why left-corner parsing metrics successfully predict brain activity of participants comprehending English (Brennan & Pylkkänen, 2017; Nelson et al., 2017). We suggested in the introduction that these effects might be weaker in languages with head-final constructions, in particular if speakers of these languages adopt predictive parsing strategies. However, in seeming conflict with this possibility, a recent study in Japanese, a strictly head-final language, showed that a left-corner parsing model outperformed a top-down parsing model in left inferior frontal and temporal-parietal regions (Sugimoto et al., 2023). One relevant difference with our study is that they used a complexity metric that considers the number of possible syntactic analyses at each word (i.e., modeling ambiguity resolution), rather than directly quantifying the number of operations that are required to build the correct structure (i.e., node count for a one-path syntactic parse tree). These metrics do not reflect the same process. It is therefore possible that brain activity corresponding to ambiguity resolution is best modeled by considering the number of syntactic analyses following a left-corner strategy, and that, when the most likely analysis is chosen, the structure-building process itself is best modeled via top-down metrics. To better understand such diverging results, it is important that future studies take into account how syntactic properties (of different languages) affect the suitability of different parsing strategies. As a case in point, it is commonly mentioned that the left-corner strategy predicts processing breakdown for exactly those constructions that are difficult to process. Left-corner parsers have the property that their memory demands increase in proportion to the number of embeddings in center-embedded constructions, while they remain constant for both right- and left- branching structures (Abney & Johnson, 1991; Johnson-Laird, 1983; Resnik, 1992). Sentences with multiple levels of center-embedding indeed quickly over-tax working memory resources (Miller & Chomsky, 1963), supporting the cognitive plausibility of the left-corner method. Intriguingly, however, processing difficulty for center-embedded constructions is not consistent across languages. Vasishth et al. (2010) found that speakers of German (a language with head-final VPs, like Dutch) are hindered less than English speakers during the comprehension of multiply center-embedded sentences, suggesting that people’s ability to generate syntactic predictions might be dependent on the specific grammatical properties of the language.

In all, the fact that node counts derived from a top-down parser best explain brain activity of people listening to Dutch stories might have to do with certain grammatical properties of Dutch, including its head-final VPs, which make left-corner prediction inadequate. That this conclusion can be reached merely by extending the approach to a closely related Germanic language underscores the need for more work on typologically diverse languages, whose structural properties invite different parsing strategies that might rely on different brain regions to varying degrees (Bornkessel-Schlesewsky & Schlesewsky, 2016). Overall, the fronto-temporal language network is remarkably consistent across speakers of different languages (Malik-Moraleda et al., 2022), but structural differences within this network can be induced by experience with sentence structures that elicit different processing behavior (Goucha et al., 2022).

### 4.2 Is node count the right linking hypothesis?

Two critical questions can be raised about our use of incremental node count as the complexity metric to represent syntax-related neural states. First, the syntactic structures that we used to compute node counts contained empty elements, such as traces (Carnie, 2021). As these do not have an acoustic correlate in the physical stimulus, we assigned their node counts to the subsequent word. We reasoned that the existence and location of an element, whether covert or overt, can usually be inferred with absolute certainty only after it has been encountered, which would be at the subsequent word. However, this wait-and-see (or wait-and-infer) attitude is somewhat inconsistent with the parsing strategy of both top-down and left-corner parsers, which build structure predictively. On the top-down method, for instance, a constituent node is postulated before there is any evidence for its existence. Given that the structure corresponding to overt elements is built predictively, it is inconsistent if the structure corresponding to covert elements is built in an integratory manner, in particular given the evidence for prediction of null forms (Lau et al., 2006; Yoshida et al., 2013).

In order to check whether the node counts for traces were assigned correctly, the reconstruction accuracy of syntactic predictors must be tested when node counts for traces are assigned to the previous word. This will shift the structural complexity of the sentence to a different point in time, and will do so differently depending on both the location of the trace and the parsing method. For traces at the left corner of a constituent (e.g., the first trace in Figure 2), whose node counts are higher for top-down than for bottom-up parsers, the preceding word will be assigned a higher node count on the top-down method. The reverse is the case for traces at the right corner of a constituent (e.g., the last trace in Figure 2), because their node counts are higher on the bottom-up method. Clearly, the method of assigning node counts of traces to the preceding or next word has important consequences, again showing that different ways of expressing syntax in the time domain lead to different predictions about the dynamics of structure building.

Another relevant question is whether incremental node count is the right measure to represent syntactic structure building. The node count metric we used is unlexicalized, which means that it does not take into account the label of the node counted. During language comprehension, however, syntactic processing is lexicalized (Coopmans et al., 2022; Hagoort, 2005), and lexical information guides predictive structure building (Arai & Keller, 2013; Boland & Blodgett, 2006; Schütze & Gibson, 1999). Such lexically driven structural predictions are represented to some extent in other metrics, such as surprisal values derived from probabilistic context-free grammars (Brennan & Hale, 2019; Brennan et al., 2016; Shain et al., 2020) or recurrent neural network grammars (Brennan et al., 2020; Hale et al., 2018; Sugimoto et al., 2023). Both types of grammars incrementally build hierarchical structure, which they use to conditionalize the probability of an upcoming word or an upcoming word’s part-of-speech. There are at least two other metrics that are informative about syntactic processing in a way that incremental node count is not. First, the “distance” metric counts the total number of syntactic analyses that are considered by a parser at every individual word (Brennan et al., 2020; Hale et al., 2018; Sugimoto et al., 2023). The larger the number of alternative analyses to be considered, the higher the effort in choosing the correct parse. In this way, it models the ambiguity resolution process that is much studied in psycholinguistics but that is not captured by node count. Second, “incremental memory” reflects the number of phrases to be held in memory on a stack (Nelson et al., 2017; Van Wagenen et al., 2014) and is sometimes quantified as the number of open nodes (Nelson et al., 2017; Giglio et al., 2024). Incremental memory is particularly relevant to evaluate left-corner parsing because one of the arguments in favor of the plausibility of the left-corner method relies on a complexity metric that reflects the number of unattached constituents to be held in working memory (Abney & Johnson, 1991). Node count instead quantifies syntactic complexity rather than memory load, and it need not be the case that these make exactly the same predictions with respect to comprehension difficulty.

## 5. Conclusions

In order for neuro-computational language models to be useful for the study of human biology, they must both be linguistically interpretable and be able to express linguistic structure in time. The current study achieves both goals by using incremental node count as the linking hypothesis that connects atemporal syntactic structure with continuously varying neural dynamics. Node counts were derived from syntactic structures that were generated via an expressive and linguistically interpretable grammar. As such, syntax was expressed in the time domain, yielding time-varying complexity metrics that could be regressed against delta-band neural activity in temporally-resolved manner. By explicitly positing which parsing strategies are mostly likely to be entertained by the brain, this approach additionally yields predictions about the functional role of these brain regions. Results show that the top-down strategy, which is maximally eager and postulates syntactic structure early in time, best reflects the neural response in inferior frontal and superior temporal regions, suggesting that these regions are involved in building syntactic structure in a predictive manner. That being said, these results stay silent on the question what kinds of computations are implemented in these areas. This has to do with the fact that we computed node count based on derived tree structures rather than on the actual derivation trees. In order to build neuro-computational language models that are closer to cognitive and neurobiological processing, it is important that future work assumes a more transparent relation between grammar and parser, for instance by using the derivation steps directly implemented by the grammar (Brennan et al., 2020; Chesi, 2015; Hale et al., 2018; Stanojević et al., 2023). Knowing when the brain projects this linguistic knowledge onto its perceptual analysis of speech is essential in order to understand how the brain transforms continuous sensory stimulation into cognitive representations, still a major question in human biology.

## Acknowledgments

We thank Laura Giglio for feedback on a previous version of this manuscript, Sophie Slaats and Michelle Suijkerbuijk for help creating the syntactic annotations, Noémie te Rietmolen for making Figure 1, and Ryan Law, Ioanna Zioga, Hugo Weissbart and Sophie Slaats for contributing to data acquisition. Peter Hagoort was supported by NWO Gravitation Grant 024.001.006 to the Language in Interaction Consortium. Andrea E. Martin was supported by an Independent Max Planck Research Group and a Lise Meitner Research Group “Language and Computation in Neural Systems”, by NWO Vidi grant 016.Vidi.188.029 to AEM, and by Big Question 5 (to Prof. dr. Roshan Cools & Dr. Andrea E. Martin) of the Language in Interaction Consortium funded by NWO Gravitation Grant 024.001.006 to PH. Cas W. Coopmans was supported by NWO Vidi grant 016.Vidi.188.029 to AEM.

## Supplementary Information

### >S1. Auditory stimuli

**Table S1.** Auditory stimuli.

| Story part | Duration |
| --- | --- |
| Andersen 1a | 4 min 58 sec |
| Andersen 1b | 5 min 17 sec |
| Andersen 1c | 4 min 49 sec |
| Andersen 1d | 5 min 50 sec |
| Grimm 1a | 6 min 6 sec |
| Grimm 1b | 6 min 40 sec |
| Grimm 2a | 5 min 3 sec |
| Grimm 2b | 5 min 32 sec |
| Grimm 2c | 5 min 2 sec |

### S2. Comparisons against the base model

The syntactic predictors are positively correlated. This is mainly the case for the left-corner predictor, whose Pearson correlation with the bottom-up and top-down predictors is 0.74 and 0.80, respectively (Figure S2.1). A high correlation between predictors in a regression analysis can lead to multicollinearity, which can in turn result in increased variance of their TRF coefficients (Weissbart et al., 2020). This issue typically emerges when the variance inflation factor (VIF) is above 5, which indicates that the variance in one predictor can be explained by a linear combination of the other predictors (Sheather, 2009). We computed the VIF for each predictor in the full model by taking the diagonal of the inverse of the correlation matrix shown in Figure S2.1. This showed that when all predictors are included, the VIF for both top-down (VIF_top-down_ = 5.09) and left-corner (VIF_left-corner_ = 9.75) is above 5, indicating that multicollinearity between the predictors might hinder TRF coefficient estimation.

**Figure S2.1.**
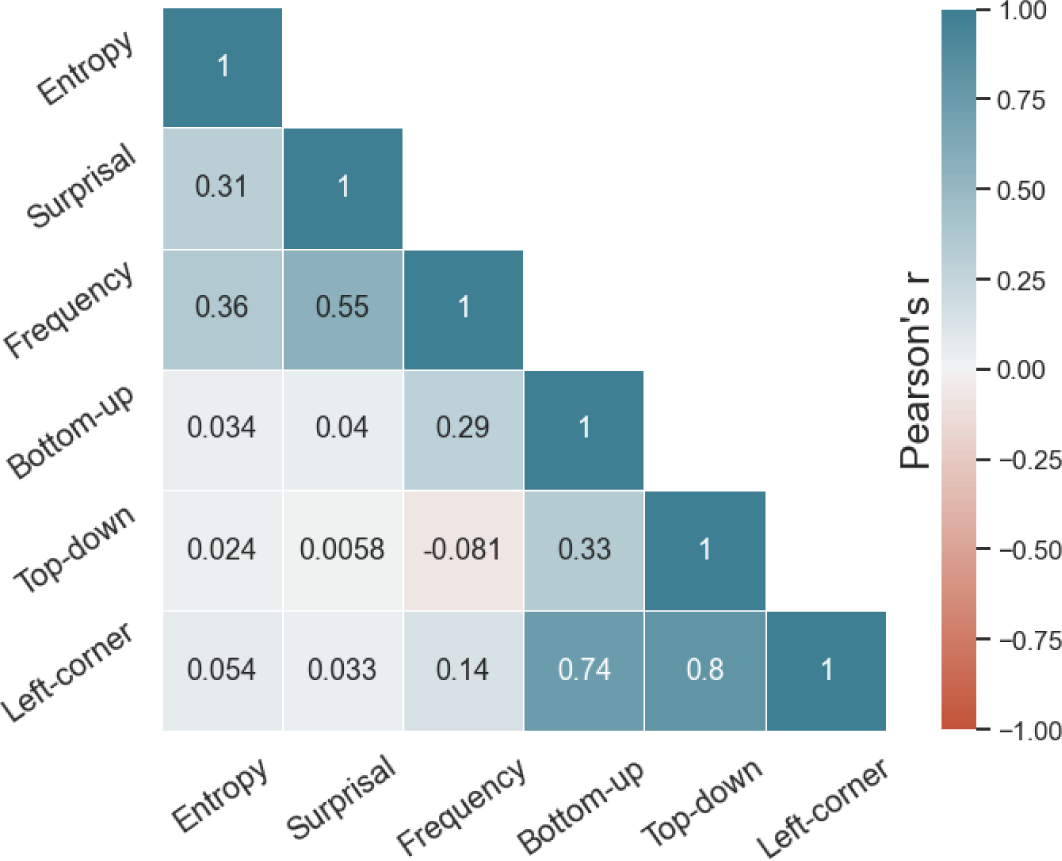
Correlation matrix showing the Pearson correlation between all word-based predictors.

As a control, we therefore repeated our analyses with TRF models in which the VIF is below 5 for all predictors. In this control analysis we evaluated whether the addition of each of the three syntactic predictors to a base model (Table S2.1) improves reconstruction accuracy. This analysis is conceptually similar to the analysis reported in the main manuscript, but it does not take into account co-dependencies between the different syntactic predictors because these predictors never appear in the same model. So, while the analyses in the main manuscript provide a conservative measure of reconstruction accuracy, the current analyses might overestimate the reconstruction accuracy of certain predictors that share variability.

**Table S2.1.** Predictors included in each model.

| Predictor(s) | Spectrogram/<br>Onsets | Word<br>onset | Word<br>frequency/<br>Entropy/<br>Surprisal | Bottom-up | Top-down | Left-corner |
| --- | --- | --- | --- | --- | --- | --- |
| **Model name** |  |  |  |  |  |  |
| **Base** | X | X | X |  |  |  |
| **Bottom-up** | X | X | X | X |  |  |
| **Top-down** | X | X | X |  | X |  |
| **Left-corner** | X | X | X |  |  | X |

#### S2.1 Model comparison

Figure S2.2A shows the sources in which the reconstruction accuracy of the base model was significantly non-zero. The spatial extent of these clusters is consistent with the location of the auditory cortex, likely reflecting the contributions of the two acoustic predictors in the base model. Figures S2.2B-D further show the sources of improvements in reconstruction accuracy when each of the three syntactic predictors are separately added to the base model. For comparison with the results reported in Section 3.1 of the main manuscript, the scales of the color bars in Figures S2.2B-D are identical to those in Figure 4. Spatial cluster-based permutation tests (cluster-level alpha = 0.0083, Bonferroni-corrected for 6 tests) revealed that the top-down predictor explains variance in frontal and temporal regions of the left hemisphere (t_av_ = 5.08, p = .0036) and to a lesser extent also the right hemisphere (t_av_ = 3.16, p < .001). The activation pattern for the left-corner predictor is rather similar, with sources of significant explained variance in the left superior temporal lobe (t_av_ = 4.19, p = .0027) and a spatially more extended area in the right frontal and temporal lobes (t_av_ = 3.09, p < .001). The bottom-up predictor engages a smaller region centered around Heschl’s gyrus, but none of these clusters are significant at the adjusted alpha level (t_av_ = 3.54, p = .0093).

**Figure S2.2.**
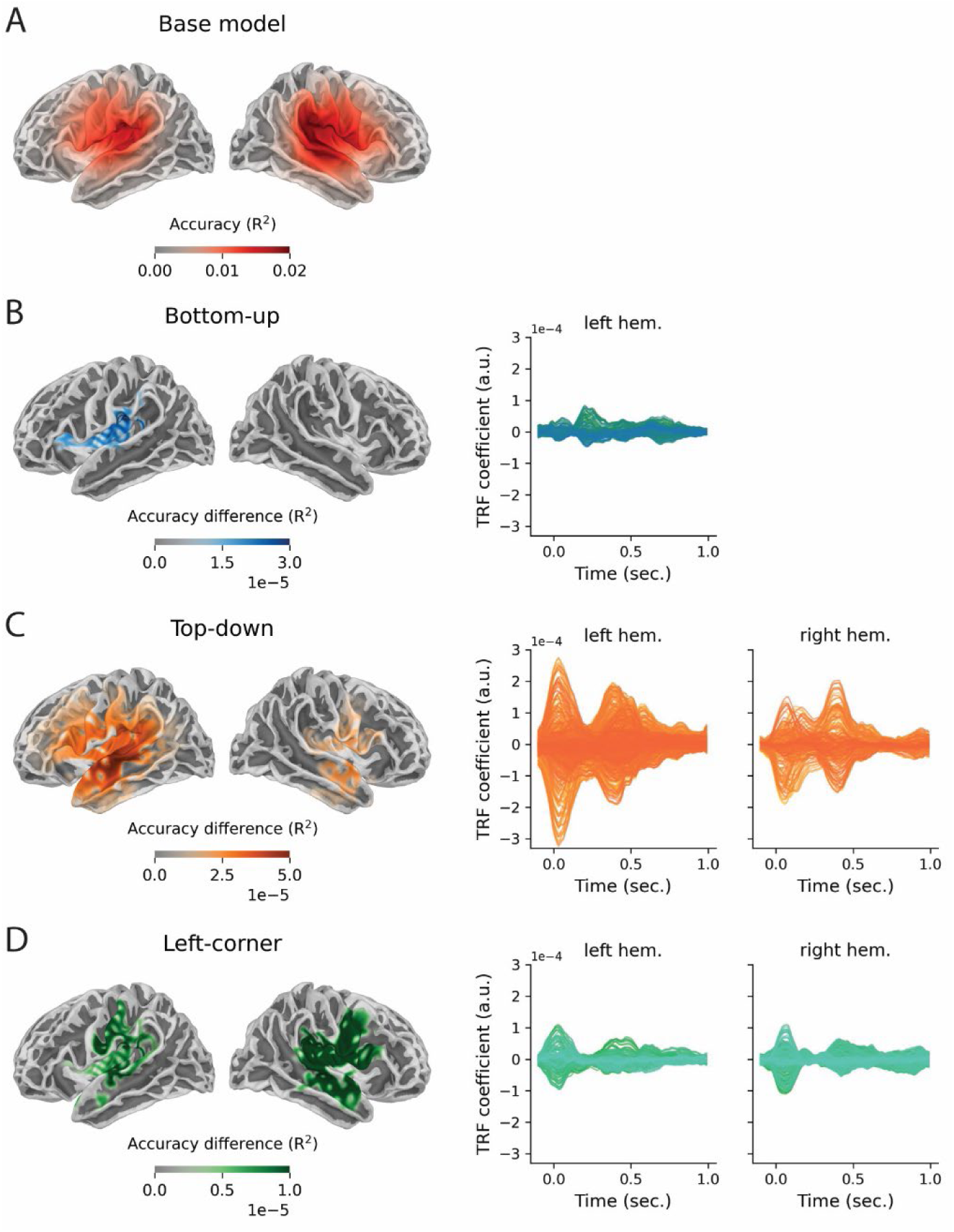
Sources of (improved) explained variance of the base model **(A)** and the predictors reflecting node count from the bottom-up **(B)**, top-down **(C)** and left-corner **(D)** methods. All clusters that were significant at uncorrected alpha = 0.05 are displayed. Notice that the scales of the color bars are different across the plots. The plots on the right show the temporal response functions for node count derived from the bottom-up, top-down, and left-corner parsers in their respective models. Each line reflects the response function in a source point that was part of a cluster which showed a significant improvement in reconstruction accuracy.

### S2.2 Evaluation of the response functions

Figure S2.2 shows the TRFs within the significant regions from the cluster-based source analysis of reconstruction accuries, in the left and the right hemisphere separately. The TRF for each syntactic predictor comes from a model which includes all base predictors as well as one syntactic predictor (e.g., the top-down TRF comes from the Top-down model; see Table S2.1). In line with the reconstruction accuracy results, the TRFs show clearly that the neural response to the information encoded in top-down node counts is stronger than the response to node counts derived from a bottom-up or a left-corner parser.

Figure S2.3 shows the source t-values (based on two-tailed, one-sample t-tests) of the TRFs for the three syntactic predictors, split up into four time windows (corresponding to delays in TRF estimation). We used spatiotemporal cluster-based permutation tests (cluster-level alpha = 0.0083, Bonferroni-corrected for 6 tests) to determine when and where the TRF coefficients of each syntactic predictor deviated from zero. Focusing on left-hemispheric sources, this analysis revealed a negative cluster for the top-down predictor in frontal and temporal regions (t_av_ = −0.45, p < .001), and a positive cluster in the middle frontal lobe (t_av_ = 0.65, p < .001). The strongest effect of the bottom-up TRF was again in a region centered around the inferior frontal cortex (t_av_ = 0.44, p < .001), and the TRF of the left-corner predictor peaked in (anterior) temporal regions (t_av_ = −0.51, p = .0073).

**Figure S2.3.**
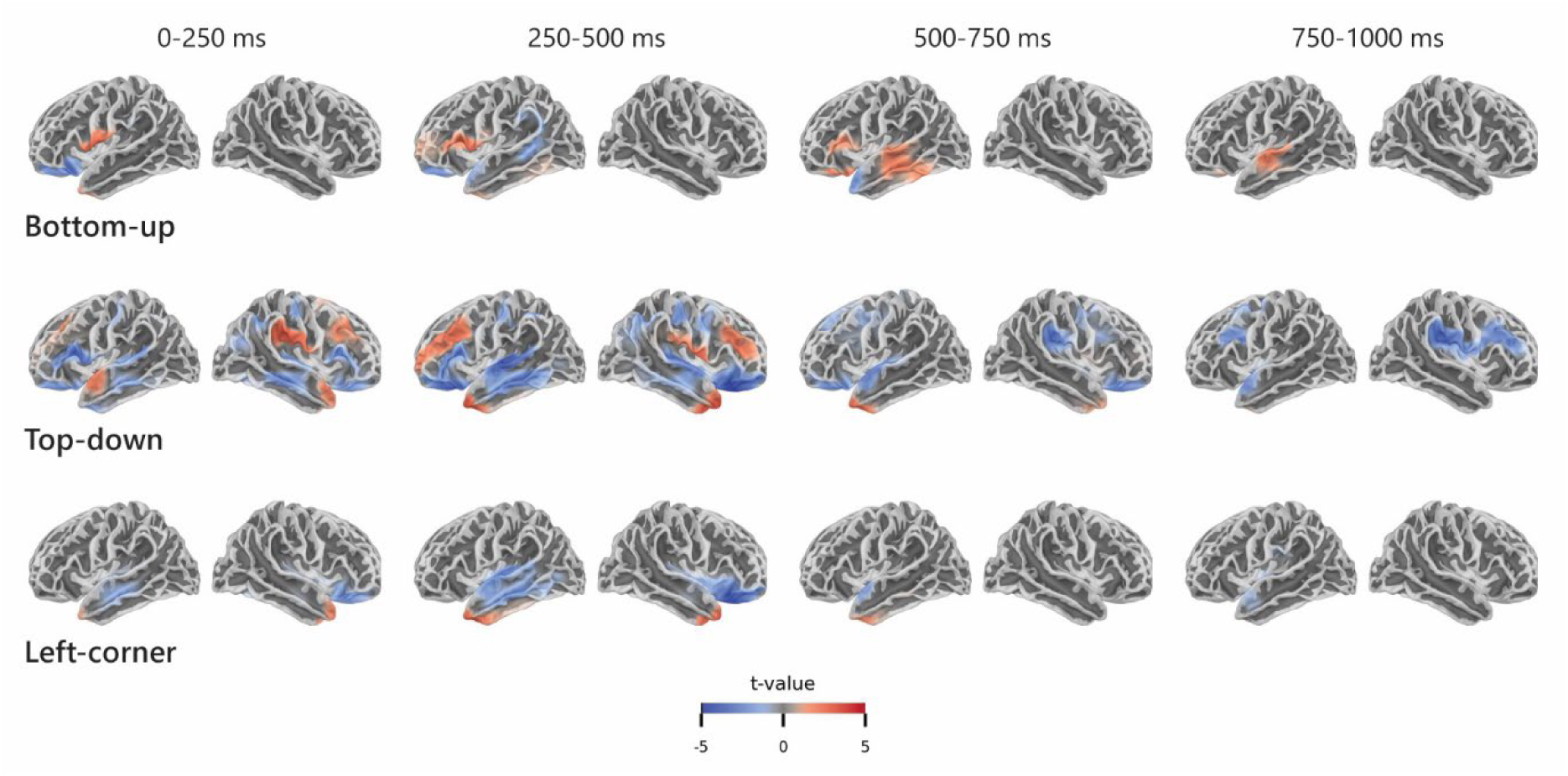
Sources of the TRFs for node count derived from bottom-up, top-down, and left-corner parsers, representing early to late responses. The colors represent positive (red) or negative (blue) t-values in sources that were significantly responsive (at corrected alpha = 0.0083) to the predictor in the indicated time windows.

### S2.3 Region of interest analysis

The following three regions of interest (ROIs) in the left hemisphere were extracted using the in the same way as described in Section 3.3 of the main manuscript: the inferior frontal gyrus (IFG), posterior temporal lobe (PTL) and anterior temporal lobe (ATL; see Figure S2.4A). Figure S2.4B shows for each ROI the improvement in reconstruction accuracy when the relevant predictor is added to the base model. Like in the main results, two-tailed paired-samples t-tests (with alpha = 0.0056, Bonferroni-corrected for 9 tests) reveal that the addition of the top-down predictor to the base model improves the reconstruction accuracy in all three ROIs (IFG: t(23) = 4.41, p < 0.001; PTL: t(23) = 5.19, p < .001; ATL: t(23) = 3.55, p = 0.0012). Adding the left-corner predictor improves reconstruction accuracy in the ATL only, t(23) = 3.34, p = 0.002.

In each of the three ROIs, we used cluster-based permutation tests to determine when the TRFs of each syntactic predictor deviated from zero. Again, this involves 9 comparisons (3 TRFs * 3 ROIs), so clusters were evaluated at alpha = 0.0056. As shown in Figure S2.5C, the top-down predictor showed effects in the IFG (from 160 to 540 ms, t_av_ = −2.88, p < .001) and PTL (from 300 to 610 ms, t_av_ = −3.01, p < .001). Quite similar to the top-down effect in the PTL, the left-corner predictor showed a significant modulation in PTL (from 330 to 480 ms, t_av_ = −3.10, p = .0042).

**Figure S2.4.**
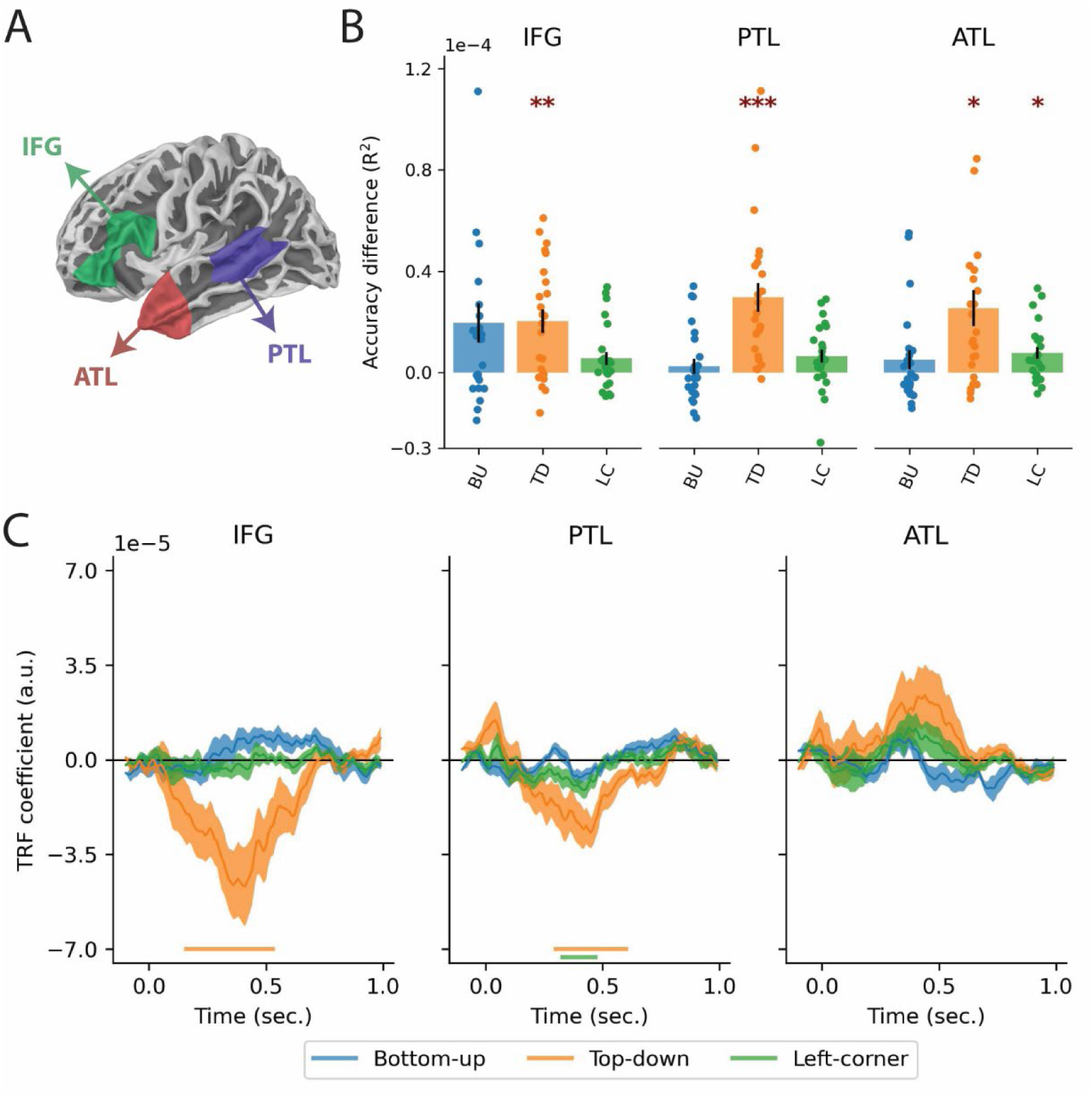
Region of interest analysis. **(A)** Spatial extensions of the three regions of interest. **(B)** Difference in reconstruction accuracy with the base model, plotted for left IFG, PTL, and ATL. The height of each bar indicates the improvement in reconstruction accuracy when only the relevant syntactic predictor was added to the base model. The drops represent the accuracy difference for individual participants, and the error bars represent the standard error of the mean across subjects. **(C)** Temporal response functions for node count derived from bottom-up, top-down, and left-corner parsers in their respective models. Error bars reflect the standard error of the mean per time sample. The horizontal bars below the TRFs reflect the time points at which the TRFs were significantly non-zero.

In all, these results are largely consistent with the results reported in the main manuscript, suggesting that multicollinearity between the predictors did not hinder estimation of the TRF coefficients. However, one notable difference has to do with the left-corner predictor, whose effects are weaker in the results reported in the main manuscript. Regarding the response functions, it is noteworthy that the TRFs for both bottom-up and top-down are stable and quite similar in the full model (Figure 6C in the main manuscript) and in the simpler models (Figure S2.4C), which shows that they are unaffected by the presence of the other syntactic predictors in the full model. The left-corner TRFs, however, are smaller in size and less variable in the full model than in the simpler Left-corner model. In the latter, the left-corner TRFs are more similar to the top-down TRFs (from the Top-down model; Figure S2.4C), suggesting that they are partially explaining the same variance. In terms of reconstruction accuracy, the explained variance in the Left-corner model attributed to the left-corner predictor (Figures S2.2D and S2.4B) is visibly reduced when the left-corner predictor is added to a null model that already contains both bottom-up and top-down as predictors (Figures 4C and 6B in the main manuscript). The reverse does not happen, suggesting that some of the variance assigned to left-corner in the simpler Left-corner model was assigned incorrectly, ‘belonging’ to top-down rather than left-corner.

### S3. The effect of predictability on integratory structure building

As an exploratory analysis of the interaction between predictability and integratory structure building, we evaluated whether the effect of bottom-up was modulated by the variable surprisal, which reflects the predictability of a word given the preceding context (Slaats et al., 2024; Tezcan et al., 2023). If demands on integration are higher for words that are not predicted, the brain response to bottom-up node counts should be larger for words that are surprising. We computed median surprisal over all stories together and then labeled each word in each story as high surprisal or low surprisal, depending on whether its surprisal was higher or lower than the overall median surprisal. However, splitting the bottom-up predictor in this way leads to two difficulties. First, such a surprisal split is confounded by word duration, because longer words are generally (less frequent and therefore) more surprising (in our stimuli: average length difference = 120 ms, t(8549) = 37.54, p < .001). We therefore iteratively sampled from the stimuli such that the high-zand low-surprisal groups were matched in word length (i.e., almost fully overlapping probability distributions whose means only differed by 0.04 ms, t(5444) = 0.01, p = 0.99). The resulting subset contains 64% of the words in original stimulus set (see Slaats et al., 2024 for discussion). A second difficulty is that splitting the bottom-up predictor by surprisal involves dichotomizing a continuous variable, which is unnatural in both neural terms (i.e., it probably does not represent how the brain transforms this kind of information) and statistical terms (i.e., it reduces power and leads to underestimated effect sizes). As a first check, we therefore compared the reconstruction accuracy of a model in which the bottom-up predictor was split into high- and low-surprisal words to the reconstruction accuracy of an equivalent model in which the bottom-up predictor was split randomly. This is an unbiased comparison because both models contain a split predictor (Slaats et al., 2024). In the random split, the two groups (bottom-up_1_ and bottom-up_2_) were selected randomly but matched in word length (average length difference = 2.03 ms, t(5444) = −0.47, p = 0.65). Reconstruction accuracy was significantly larger for the model in which the bottom-up predictor was split by surprisal than for the model in which it was split randomly (largest cluster t_av_ = 4.01, p < .001). This provides a first indication that splitting bottom-up node counts by their surprisal value captures something relevant; it suggests that predictability modulates the neural response to bottom-up structure building.

To further explore this interaction, we evaluated the effects of bottom-up for high- vs. low-surprisal words. This analysis involved comparing the reconstruction accuracy of a model with bottom-up for only high-surprisal words and a model with bottom-up for only low-surprisal words. Both models contained all other predictors and only differed in whether the bottom-up predictor reflected high- or low-surprisal words. These models did differ in reconstruction accuracy, but not as expected: significantly more variance was explained by the model containing the bottom-up predictor for low-surprisal words than the model containing the bottom-up predictor for high-surprisal words (largest cluster t_av_ = 3.59, p < .001; Figure S3.1A). Likewise, the temporal response function of bottom-up for low-surprisal words is more pronounced (Figures S3.1B and S3.1C). This suggests, contrary to our hypothesis, that the brain’s response to the (syntactic properties of the) bottom-up input is stronger when the input can be predicted (for related results, and a more comprehensive exploration of the relation between syntactic and distributional information, see Slaats et al., 2024).

**Figure S3.1.**
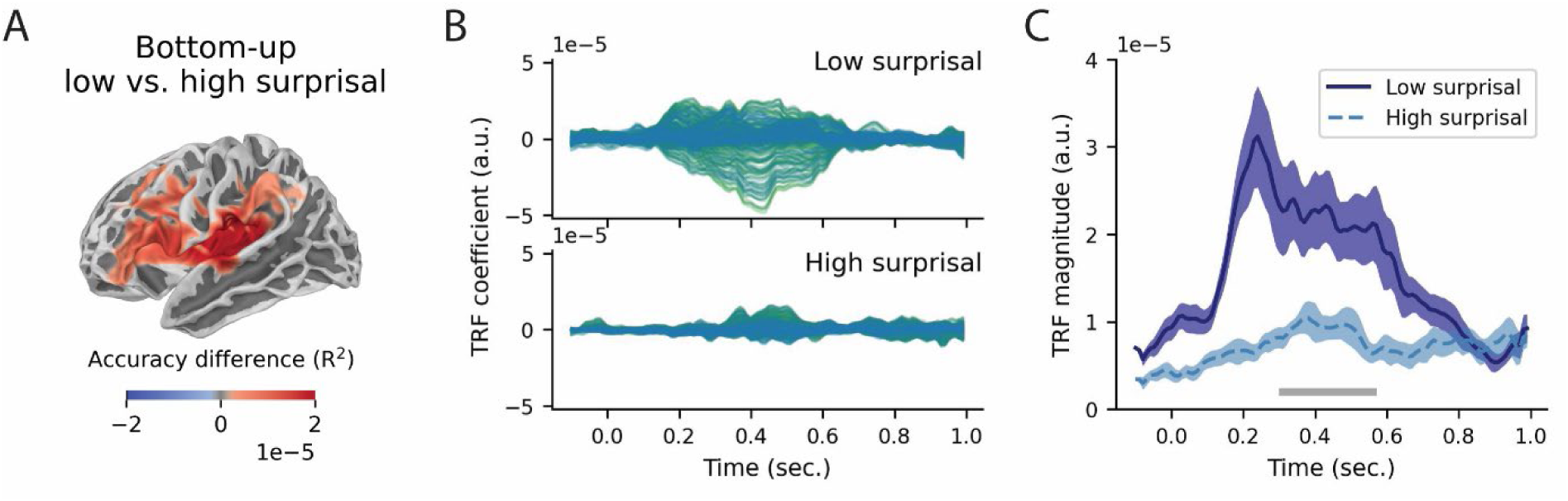
The effect of predictability on integratory structure building. **(A)** Significant sources of the accuracy differences between bottom-up node counts for low- vs. high-surprisal words. The positive accuracy difference indicates that the reconstruction accuracy was higher for the model with low- than for the model with high-surprisal words. **(B)** Temporal response functions for the bottom-up predictor, estimated separately for high-surprisal and low-surprisal words. Each line reflects the TRF estimated for a source point that was significant in the reconstruction accuracy analysis (i.e., the colored areas in (A)). **(C)** Amplitude of the response functions, averaged over significant sources. The error bars reflect the standard error of the mean per time sample. The horizontal bar below the TRFs reflects the temporal extension of the largest cluster indicating the significant difference between the TFR amplitudes.

It thus appears to be the case that predictability modulates integratory structure building, but the story is not as straightforward as sketched in the main manuscript, at least not when a general measure of predictability like surprisal is used. That is, we used *lexical* surprisal (i.e., surprisal about the current word) rather than *structural* surprisal (i.e., surprisal about the syntactic analysis demanded by the current word), which likely affects the interaction with bottom-up integration cost. When structural surprisal is high, the predicted syntactic analysis is wrong, yielding high demands on bottom-up structural integration. In contrast, when lexical surprisal is high, it simply means that the current word was not predicted. But that does not necessarily mean that syntactic integration is costly, as it might very well be that the word’s part of speech was predicted correctly, due to which integrating that (surprising) word into the syntactic structure is relatively straightforward. Another complicating factor is that the surprisal split is confounded not only with word duration, but also with word frequency (because surprisal is correlated with frequency, see Figure S2.1) and to a lesser extent also with entropy and the position of a word in a sentence (i.e., words in early-sentence positions are generally more surprising). All of these correlations might affect the pattern we see here, which underscores the importance of incorporating interactive effects in encoding models of language-related brain activity (see Slaats et al., 2024 for an exploration).

## Footnotes

1 Note that the notions ‘top’ and ‘bottom’ in this context refer to the geometry of syntactic tree structures. Top-down thus says something about the (vertical) direction of phrase structure building, and with ‘top-down effects’ we refer to the effects of node count derived by a top-down parser. In the predictive processing literature, the term ‘top-down effects’ is commonly used to describe how low-level processes are affected by high-level sources of information, but this is not our intended interpretation.

1 These average cluster-level t-values are underestimated because they are corrected for the total number of significant source points across the whole duration of the cluster. As not all of these source points are significantly involved at all time points (and might at certain time points even have a sign opposing the sign of the cluster), the correction is overly strict.

## References

Abney, S. P., & Johnson, M. (1991). Memory requirements and local ambiguities of parsing strategies. Journal of Psycholinguistic Research, 20(3), 233–250. 10.1007/BF01067217

Agmon, G., Jaeger, M., Tsarfaty, R., Bleichner, M. G., & Zion Golumbic, E. (2023). “Um…, It’s Really Difficult to… Um… Speak Fluently”: Neural Tracking of Spontaneous Speech. Neurobiology of Language, 4(3), 435–454. 10.1162/nol_a_00109

Arai, M., & Keller, F. (2013). The use of verb-specific information for prediction in sentence processing. Language and Cognitive Processes, 28(4), 525–560. 10.1080/01690965.2012.658072

Bai, F., Meyer, A. S., & Martin, A. E. (2022). Neural dynamics differentially encode phrases and sentences during spoken language comprehension. PLOS Biology, 20(7), e3001713. 10.1371/journal.pbio.3001713

Bhattasali, S., Fabre, M., Luh, W.-M., Saied, H. A., Constant, M., Pallier, C., Brennan, J. R., Spreng, R. N., & Hale, J. (2019). Localising memory retrieval and syntactic composition: An fMRI study of naturalistic language comprehension. *Language*, Cognition and Neuroscience, 34(4), 491–510. 10.1080/23273798.2018.1518533

Boland, J. E., & Blodgett, A. (2006). Argument Status and PP-Attachment. Journal of Psycholinguistic Research, 35(5), 385–403. 10.1007/s10936-006-9021-z

Bornkessel-Schlesewsky, I., & Schlesewsky, M. (2016). The importance of linguistic typology for the neurobiology of language. Linguistic Typology, 20(3), 241–252. 10.1515/lingty-2016-0032

Brainard, D. H. (1997). The Psychophysics Toolbox. Spatial Vision, 10(4), 433–436.

Brennan, J. (2016). Naturalistic Sentence Comprehension in the Brain. Language and Linguistics Compass, 10(7), 299–313. 10.1111/lnc3.12198

Brennan, J. R., Dyer, C., Kuncoro, A., & Hale, J. T. (2020). Localizing syntactic predictions using recurrent neural network grammars. Neuropsychologia, 146, 107479. 10.1016/j.neuropsychologia.2020.107479

Brennan, J. R., & Hale, J. T. (2019). Hierarchical structure guides rapid linguistic predictions during naturalistic listening. PLOS ONE, 14(1), e0207741. 10.1371/journal.pone.0207741

Brennan, J. R., & Martin, A. E. (2020). Phase synchronization varies systematically with linguistic structure composition. Philosophical Transactions of the Royal Society B: Biological Sciences, 375(1791), 20190305. 10.1098/rstb.2019.0305

Brennan, J. R., & Pylkkänen, L. (2017). MEG Evidence for Incremental Sentence Composition in the Anterior Temporal Lobe. Cognitive Science, 41(S6), 1515–1531. 10.1111/cogs.12445

Brennan, J. R., Stabler, E. P., Van Wagenen, S. E., Luh, W.-M., & Hale, J. T. (2016). Abstract linguistic structure correlates with temporal activity during naturalistic comprehension. Brain and Language, 157–158, 81–94. 10.1016/j.bandl.2016.04.008

Brennan, J., Nir, Y., Hasson, U., Malach, R., Heeger, D. J., & Pylkkänen, L. (2012). Syntactic structure building in the anterior temporal lobe during natural story listening. Brain and Language, 120(2), 163–173. 10.1016/j.bandl.2010.04.002

Brodbeck, C., Jiao, A., Hong, L. E., & Simon, J. Z. (2020). Neural speech restoration at the cocktail party: Auditory cortex recovers masked speech of both attended and ignored speakers. PLOS Biology, 18(10), e3000883. 10.1371/journal.pbio.3000883

Brodbeck, C., Bhattasali, S., Cruz Heredia, A. A., Resnik, P., Simon, J. Z., & Lau, E. (2022). Parallel processing in speech perception with local and global representations of linguistic context. ELife, 11, e72056. 10.7554/eLife.72056

Brodbeck, C., Brooks, T. L., Das, P., Reddigari, S., & Kulasingham, J. P. (2021). Eelbrain 0.37. Zenodo. 10.5281/zenodo.4650416

Brodbeck, C., Presacco, A., & Simon, J. Z. (2018). Neural source dynamics of brain responses to continuous stimuli: Speech processing from acoustics to comprehension. NeuroImage, 172, 162–174. 10.1016/j.neuroimage.2018.01.042

Carnie, A. (2021). Syntax: A Generative Introduction. Wiley-Blackwell.

Chesi, C. (2015). On directionality of phrase structure building. Journal of Psycholinguistic Research, 44(1), 65–89. 10.1007/s10936-014-9330-6

Coopmans, C. W., & Schoenmakers, G.-J. (2020). Incremental structure building of preverbal PPs in Dutch. Linguistics in the Netherlands, 37, 38–52. 10.1075/avt.00036.coo

Coopmans, C. W., de Hoop, H., Hagoort, P., & Martin, A. E. (2022). Effects of Structure and Meaning on Cortical Tracking of Linguistic Units in Naturalistic Speech. Neurobiology of Language, 3(3), 386–412. 10.1162/nol_a_00070

Coopmans, C. W., & Zaccarella, E. (2023). Three conceptual clarifications about syntax and the brain. Frontiers in Language Sciences, 2. 10.3389/flang.2023.1218123

Dale, A. M., Fischl, B., & Sereno, M. I. (1999). Cortical Surface-Based Analysis: I. Segmentation and Surface Reconstruction. NeuroImage, 9(2), 179–194. 10.1006/nimg.1998.0395

Daube, C., Ince, R. A. A., & Gross, J. (2019). Simple Acoustic Features Can Explain Phoneme-Based Predictions of Cortical Responses to Speech. Current Biology, 29(12), 1924–1937.e9. 10.1016/j.cub.2019.04.067

David, S. V., Mesgarani, N., & Shamma, S. A. (2007). Estimating sparse spectro-temporal receptive fields with natural stimuli. Network: Computation in Neural Systems, 18(3), 191–212. 10.1080/09548980701609235

de Vries, W., & Nissim, M. (2021). As Good as New. How to Successfully Recycle English GPT-2 to Make Models for Other Languages. Findings of the Association for Computational Linguistics: ACL-IJCNLP 2021, 836–846. 10.18653/v1/2021.findings-acl.74

Demberg, V., & Keller, F. (2019). Cognitive models of syntax and sentence processing. In P. Hagoort (Ed.), Human Language: From Genes and Brains to Behavior (pp. 293–312). MIT Press.

Ding, N., Melloni, L., Zhang, H., Tian, X., & Poeppel, D. (2016). Cortical tracking of hierarchical linguistic structures in connected speech. Nature Neuroscience, 19(1), 158–164. 10.1038/nn.4186

Embick, D., & Poeppel, D. (2015). Towards a computational(ist) neurobiology of language: Correlational, integrated and explanatory neurolinguistics. Language, Cognition and Neuroscience, 30(4), 357–366. 10.1080/23273798.2014.980750

Ferreira, F., & Qiu, Z. (2021). Predicting syntactic structure. Brain Research, 1770, 147632. 10.1016/j.brainres.2021.147632

Frazier, L. (1985). Syntactic complexity. In A. M. Zwicky, D. R. Dowty, & L. Karttunen (Eds.), Natural Language Parsing: Psychological, Computational, and Theoretical Perspectives (pp. 129–189). Cambridge University Press.

Giglio, L., Sharoh, D., Ostarek, M., & Hagoort, P. (2024). Diverging neural dynamics for syntactic structure building in naturalistic speaking and listening. Proceedings of the National Academy of Sciences, 121(11), e2310766121. 10.1073/pnas.2310766121

Giglio, L., Ostarek, M., Weber, K., & Hagoort, P. (2022). Commonalities and Asymmetries in the Neurobiological Infrastructure for Language Production and Comprehension. Cerebral Cortex, 32(7), 1405–1418. 10.1093/cercor/bhab287

Gillis, M., Vanthornhout, J., Simon, J. Z., Francart, T., & Brodbeck, C. (2021). Neural Markers of Speech Comprehension: Measuring EEG Tracking of Linguistic Speech Representations, Controlling the Speech Acoustics. Journal of Neuroscience, 41(50), 10316–10329. 10.1523/JNEUROSCI.0812-21.2021

Goucha, T., Anwander, A., Adamson, H., & Friederici, A. D. (2022). Native language leaves distinctive traces in brain connections. bioRxiv. 10.1101/2022.07.30.501987

Hagoort, P. (2005). On Broca, brain, and binding: A new framework. Trends in Cognitive Sciences, 9(9), 416–423. 10.1016/j.tics.2005.07.004

Hagoort, P., & Indefrey, P. (2014). The neurobiology of language beyond single words. Annual Review of Neuroscience, 37(1), 347–362. 10.1146/annurev-neuro-071013-013847

Hale, J. (2001). A probabilistic early parser as a psycholinguistic model. In Proceedings of the Second Meeting of the North American Chapter of the Association for Computational Linguistics on Language Technologies (pp. 1–8). 10.3115/1073336.1073357

Hale, J. T. (2014). Automaton theories of human sentence comprehension. CSLI Publications.

Hale, J. T., Campanelli, L., Li, J., Bhattasali, S., Pallier, C., & Brennan, J. R. (2022). Neurocomputational Models of Language Processing. Annual Review of Linguistics, 8(1), 427–446. 10.1146/annurev-linguistics-051421-020803

Hale, J., Dyer, C., Kuncoro, A., & Brennan, J. (2018). Finding syntax in human encephalography with beam search. Proceedings of the 56th Annual Meeting of the Association for Computational Linguistics (Volume 1: Long Papers), 2727–2736. 10.18653/v1/P18-1254

Hartwigsen, G. (2018). Flexible Redistribution in Cognitive Networks. Trends in Cognitive Sciences, 22(8), 687–698. 10.1016/j.tics.2018.05.008

Heeris, J. (2018). Gammatone Filterbank Toolkit. Github. https://github.com/detly/gammatone

Heilbron, M., Armeni, K., Schoffelen, J.-M., Hagoort, P., & de Lange, F. P. (2022). A hierarchy of linguistic predictions during natural language comprehension. Proceedings of the National Academy of Sciences, 119(32), e2201968119. 10.1073/pnas.2201968119

Henke, L., & Meyer, L. (2021). Endogenous oscillations time-constrain linguistic segmentation: Cycling the garden path. Cerebral Cortex, 31(9), 4289–4299. 10.1093/cercor/bhab086

Hultén, A., Schoffelen, J.-M., Uddén, J., Lam, N. H. L., & Hagoort, P. (2019). How the brain makes sense beyond the processing of single words – An MEG study. NeuroImage, 186, 586–594. 10.1016/j.neuroimage.2018.11.035

Jackendoff, R. (1977). X’ Syntax: A Theory of Phrase Structure. MIT Press. Johnson-Laird, P. N. (1983). Mental Models. Harvard University Press.

Kaufeld, G., Bosker, H. R., ten Oever, S., Alday, P. M., Meyer, A. S., & Martin, A. E. (2020). Linguistic Structure and Meaning Organize Neural Oscillations into a Content-Specific Hierarchy. The Journal of Neuroscience, 40(49), 9467–9475. 10.1523/JNEUROSCI.0302-20.2020

Keuleers, E., Brysbaert, M., & New, B. (2010). SUBTLEX-NL: A new measure for Dutch word frequency based on film subtitles. Behavior Research Methods, 42(3), 643–650. 10.3758/BRM.42.3.643

Kisler, T., Reichel, U., & Schiel, F. (2017). Multilingual processing of speech via web services. Computer Speech & Language, 45, 326–347. 10.1016/j.csl.2017.01.005

Kuperberg, G. R., & Jaeger, T. F. (2016). What do we mean by prediction in language comprehension? Language, Cognition and Neuroscience, 31(1), 32–59. 10.1080/23273798.2015.1102299

Lau, E., Stroud, C., Plesch, S., & Phillips, C. (2006). The role of structural prediction in rapid syntactic analysis. Brain and Language, 98(1), 74–88. 10.1016/j.bandl.2006.02.003

Li, J., & Hale, J. (2019). Grammatical predictors for fMRI time-courses. In R. C. Berwick & E. P. Stabler (Eds.), Minimalist Parsing (pp. 159–173). Oxford University Press. 10.1093/oso/9780198795087.003.0007

Lopopolo, A., van den Bosch, A., Petersson, K.-M., & Willems, R. M. (2021). Distinguishing Syntactic Operations in the Brain: Dependency and Phrase-Structure Parsing. Neurobiology of Language, 2(1), 152–175. 10.1162/nol_a_00029

Malik-Moraleda, S., Ayyash, D., Gallée, J., Affourtit, J., Hoffmann, M., Mineroff, Z., Jouravlev, O., & Fedorenko, E. (2022). An investigation across 45 languages and 12 language families reveals a universal language network. Nature Neuroscience, 25(8), 1014–1019. 10.1038/s41593-022-01114-5

Maris, E., & Oostenveld, R. (2007). Nonparametric statistical testing of EEG- and MEG-data. Journal of Neuroscience Methods, 164(1), 177–190. 10.1016/j.jneumeth.2007.03.024

Martin, A. E. (2016). Language processing as cue integration: Grounding the psychology of language in perception and neurophysiology. Frontiers in Psychology, 7. 10.3389/fpsyg.2016.00120

Martin, A. E. (2020). A compositional neural architecture for language. Journal of Cognitive Neuroscience, 32(8), 1407–1427. 10.1162/jocn_a_01552

Martin, A. E., & Doumas, L. A. A. (2017). A mechanism for the cortical computation of hierarchical linguistic structure. PLOS Biology, 15(3), e2000663. 10.1371/journal.pbio.2000663

Martin, A. E., & Doumas, L. A. (2019). Predicate learning in neural systems: using oscillations to discover latent structure. Current Opinion in Behavioral Sciences, 29, 77–83.

Matar, S., Dirani, J., Marantz, A., & Pylkkänen, L. (2021). Left posterior temporal cortex is sensitive to syntax within conceptually matched Arabic expressions. Scientific Reports, 11(1), 7181. 10.1038/s41598-021-86474-x

Matchin, W., Brodbeck, C., Hammerly, C., & Lau, E. (2019). The temporal dynamics of structure and content in sentence comprehension: Evidence from fMRI-constrained MEG. Human Brain Mapping, 40(2), 663–678. 10.1002/hbm.24403

Matchin, W., Hammerly, C., & Lau, E. (2017). The role of the IFG and pSTS in syntactic prediction: Evidence from a parametric study of hierarchical structure in fMRI. Cortex, 88, 106–123. 10.1016/j.cortex.2016.12.010

Meyer, L., Sun, Y., & Martin, A. E. (2020). Synchronous, but not entrained: Exogenous and endogenous cortical rhythms of speech and language processing. Language, Cognition and Neuroscience, 35(9), 1089–1099. 10.1080/23273798.2019.1693050

Miller, G. A., & Chomsky, N. (1963). Finitary models of language users. In R. D. Luce, R. R. Bush, & E. Galanter (Eds.), Handbook of Mathematical Psychology II (pp. 419–491). Wiley.

Nelson, M. J., Karoui, I. E., Giber, K., Yang, X., Cohen, L., Koopman, H., Cash, S. S., Naccache, L., Hale, J. T., Pallier, C., & Dehaene, S. (2017). Neurophysiological dynamics of phrase-structure building during sentence processing. Proceedings of the National Academy of Sciences, 114(18), E3669–E3678. 10.1073/pnas.1701590114

Pallier, C., Devauchelle, A.-D., & Dehaene, S. (2011). Cortical representation of the constituent structure of sentences. Proceedings of the National Academy of Sciences, 108(6), 2522–2527. 10.1073/pnas.1018711108

Poeppel, D. (2012). The maps problem and the mapping problem: Two challenges for a cognitive neuroscience of speech and language. Cognitive Neuropsychology, 29(1–2), 34–55. 10.1080/02643294.2012.710600

Resnik, P. (1992). Left-corner parsing and psychological plausibility. In Proceedings of the 14th International Conference on Computational Linguistics (pp. 191–197).

Schütze, C. T., & Gibson, E. (1999). Argumenthood and English prepositional phrase attachment. Journal of Memory and Language, 40(3), 409–431. 10.1006/jmla.1998.2619

Shain, C., Blank, I. A., van Schijndel, M., Schuler, W., & Fedorenko, E. (2020). FMRI reveals language-specific predictive coding during naturalistic sentence comprehension. Neuropsychologia, 138, 107307. 10.1016/j.neuropsychologia.2019.107307

Sheather, S. J. (2009). Diagnostics and transformations for multiple linear regression. In S. Sheather (Ed.), A modern approach to regression with R (pp. 151–225). Springer. 10.1007/978-0-387-09608-7_6

Slaats, S., Meyer, A., & Martin, A. E. (2024). Lexical surprisal shapes the time course of syntactic structure building. PsyArXiv. 10.31219/osf.io/fd8mh

Snijders, T. M., Vosse, T., Kempen, G., Van Berkum, J. J. A., Petersson, K. M., & Hagoort, P. (2009). Retrieval and Unification of Syntactic Structure in Sentence Comprehension: An fMRI Study Using Word-Category Ambiguity. Cerebral Cortex, 19(7), 1493–1503. 10.1093/cercor/bhn187

Sprouse, J., & Hornstein, N. (2016). Syntax and the cognitive neuroscience of syntactic structure building. In G. Hickok & S. L. Small (Eds.), Neurobiology of language (pp. 165–174). Elsevier.

Stanojević, M., Brennan, J. R., Dunagan, D., Steedman, M., & Hale, J. T. (2023). Modeling Structure-Building in the Brain With CCG Parsing and Large Language Models. Cognitive Science, 47(7), e13312. 10.1111/cogs.13312

Staub, A., & Clifton, C. (2006). Syntactic prediction in language comprehension: Evidence from either…or. *Journal of Experimental Psychology: Learning*, Memory, and Cognition, 32(2), 425. 10.1037/0278-7393.32.2.425

Stolk, A., Todorovic, A., Schoffelen, J.-M., & Oostenveld, R. (2013). Online and offline tools for head movement compensation in MEG. NeuroImage, 68, 39–48. 10.1016/j.neuroimage.2012.11.047

Sugimoto, Y., Yoshida, R., Jeong, H., Koizumi, M., Brennan, J. R., & Oseki, Y. (2023). Localizing Syntactic Composition with Left-Corner Recurrent Neural Network Grammars. Neurobiology of Language, 1–24. 10.1162/nol_a_00118

ten Oever, S., Carta, S., Kaufeld, G., & Martin, A. E. (2022). Neural tracking of phrases in spoken language comprehension is automatic and task-dependent. ELife, 11, e77468. 10.7554/eLife.77468

Tezcan, F., Weissbart, H., & Martin, A. E. (2023). A tradeoff between acoustic and linguistic feature encoding in spoken language comprehension. eLife, 12, e82386. 10.7554/eLife.82386

Van Wagenen, S., Brennan, J., & Stabler, E. P. (2014). Quantifying parsing complexity as a function of grammar. UCLA Working Papers in Linguistics, 18, 31–47.

Vasishth, S., Suckow, K., Lewis, R. L., & Kern, S. (2010). Short-term forgetting in sentence comprehension: Crosslinguistic evidence from verb-final structures. Language and Cognitive Processes, 25(4), 533–567. 10.1080/01690960903310587

Wei, K., & Körding, K. P. (2011). Causal Inference in Sensorimotor Learning and Control. In J. Trommershäuser, K. Kording, & M. S. Landy (Eds.), Sensory Cue Integration (p. 30–45). Oxford University Press. 10.1093/acprof:oso/9780195387247.003.0002

Weissbart, H., Kandylaki, K. D., & Reichenbach, T. (2020). Cortical tracking of surprisal during continuous speech comprehension. Journal of Cognitive Neuroscience, 32(1), 155–166. 10.1162/jocn_a_01467

Yoshida, M., Dickey, M. W., & Sturt, P. (2013). Predictive processing of syntactic structure: Sluicing and ellipsis in real-time sentence processing. Language and Cognitive Processes, 28(3), 272–302. 10.1080/01690965.2011.622905

Zaccarella, E., Meyer, L., Makuuchi, M., & Friederici, A. D. (2017). Building by syntax: The neural basis of minimal linguistic structures. Cerebral Cortex, 27(1), 411–421. 10.1093/cercor/bhv234

